# Integrated NMR and cryo-EM atomic-resolution structure determination of a half-megadalton enzyme complex

**DOI:** 10.1101/498287

**Authors:** Diego F. Gauto, Leandro F. Estrozi, Charles D. Schwieters, Gregory Effantin, Pavel Macek, Remy Sounier, Astrid C. Sivertsen, Elena Schmidt, Rime Kerfah, Guillaume Mas, Jacques-Philippe Colletier, Peter Güntert, Adrien Favier, Guy Schoehn, Jerome Boisbouvier, Paul Schanda

## Abstract

Atomic-resolution structure determination is the key requirement for understanding protein function. Cryo-EM and NMR spectroscopy both provide structural information, but currently cryo-EM does not routinely give access to atomic-level structural data, and, generally, NMR structure determination is restricted to small (<30 kDa) proteins. We introduce an integrated structure determination approach that simultaneously uses NMR and EM data to overcome the limits of each of these methods. The approach enabled determination of the high-resolution structure of the 468 kDa large dodecameric aminopeptidase TET2 to a precision and accuracy below 1 Å by combining secondary-structure information obtained from near-complete magic-angle-spinning NMR assignments of the 39 kDa-large subunits, distance restraints from backbone amides and specifically labelled methyl groups, and a 4.1 Å resolution EM map. The resulting structure exceeds current standards of NMR and EM structure determination in terms of molecular weight and precision. Importantly, the approach is successful even in cases where only medium-resolution (up to 8 Å) cryo-EM data are available, thus paving avenues for the structure determination of challenging biological assemblies.

---

For decades, the weight of atomic-resolution structure elucidation, a requirement for understanding biomolecular mechanisms in detail, has been almost exclusively borne by X-ray crystallography, as highlighted by the fact that about 90 % of all entries in the Protein Data Bank (PDB) are crystal structures. Notwithstanding this success of crystallography, many supra-molecular edifices, self-assembling systems, membrane proteins and proteins with extended dynamic domains are difficult to crystallize, or the crystals do not diffract to high resolution. Single-particle cryo electron microscopy (cryo-EM) and nuclear magnetic resonance spectroscopy (NMR) are not bound to obtaining well-ordered crystals, and are applicable even in the presence of significant motions. Enabled by decisive instrumental and methodological progress, cryo-EM has recently made a leap in resolution ^1,2^, which opened avenues for determining protein structures at atomic detail. Nonetheless, *de novo* atomic-resolution structure determination from cryo-EM data is currently not the rule. A survey of the electron density maps deposited in the EM data base (EMDB) over the last two years reveals an average resolution in the range of 6.5-8 Å. In 2016, a particularly productive year for single-particle cryo-EM (961 entries in the EMDB), only 20% of the deposited maps had a resolution of 3.5 Å or better, thus being well suited for atomic-level de novo structure determination. Extending EM structure determination to a wider range of biological objects may require further resolution increase and/or combination of EM with other data.

NMR spectroscopy probes the structure of proteins on a length scale that is very different from that seen by EM. Rather than probing the electronic potential of the molecule, providing a more or less well-defined molecular envelope, NMR detects the immediate vicinity of hundreds to thousands of individual atomic nuclei across the protein. Their resonance frequency is exquisitely sensitive to the local electronic environment and, thus, to the local structure. Furthermore, dedicated NMR experiments identify which atoms are in the vicinity (< 5-10 Å) of a given atomic nucleus, thus allowing to assemble structural elements into a full three-dimensional structure. The precision and accuracy of NMR-derived structures strongly depend on the availability of a large number of such distance restraints across the entire molecule. When NMR structure determination fails, it is generally the paucity of meaningful distance restraints – often as a consequence of resonance overlap or low detection sensitivity – which hampers the arrangement of local structural elements to a well-defined tertiary fold. In solution-state NMR, structure determination is particularly challenging for proteins larger than 50 kDa, as the slow molecular tumbling leads to rapid nuclear relaxation and thereby low detection sensitivity, which results in difficulties obtaining information about local conformation and distances. Consequently, besides three cases^3–5^ the PDB is, to our knowledge, devoid of *de novo* NMR structures beyond 50 kDa (subunit size). The introduction of deuteration and specific methyl labeling with ^13^CH_3_ groups has greatly expanded the molecular weight range in which solution-NMR signals can be detected to ca. 1 MDa.^6–8^ Yet because this approach is limited to methyl groups, no information about the backbone conformation can be obtained, which excludes any *de novo* structure determination.

Magic-angle spinning (MAS) solid-state NMR, overcomes the inherent size limitations that its solution-state counterpart faces by replacing the slow stochastic molecular tumbling in solution – which is at the origin of the rapid NMR signal loss – by an immobilization of the protein, and spinning of the sample at a constant “magic” angle.^9^ Recent advances in MAS NMR sample preparation, including optimized isotope-labeling schemes, hardware allowing fast sample spinning (>50 kHz), and optimized pulse sequences have enabled the structure determination of small to medium-sized proteins forming crystals, oligomeric assemblies, or embedded in membranes, generally with monomer sizes below 20 kDa,^10^ with an exceptional case at 27 kDa.^11^ Sensitivity limitations and the increased spectral complexity and signal overlap in large proteins have so far been a bottleneck for the extraction of sufficient useful restraints for structure determination on larger proteins.

While EM and NMR spectroscopy probe protein structure on different length scales, these complementary views may be combined. Recent work has explored the joint use of NMR and EM data to determine protein structures.^12–14^ The proteins studied are small (< 100 residues) and comprise only 2 or 3 secondary structure elements, which made it possible to either first determine a global fold from NMR and refine this model with the aid of the EM map^12^, or to build a first model into the electron density map, and use NMR data to refine these manually constructed EM models^13,14^. For larger proteins, the situation is significantly more complex: the former approach, i. e., determining a rough fold from NMR and then refining with EM data, is challenging because lack of NMR restraints—often the case for large proteins—precludes obtaining a converged structural model which then could be used for EM-based refinement. The latter method suffers from the fact that *de novo* building of the protein sequence into an electron density, i.e., unambiguously identifying the residues and being able to follow the chain throughout the electron density, becomes increasingly complex with protein size, and very strongly depends on the resolution of the EM data.

Here, we present an approach that overcomes these challenges and combines NMR and EM information in a joint and lightly-supervised manner. Briefly, the key ingredients for the combined analysis from these two techniques are (i) the identification of structural features (such as *α*-helices) in 3D space from EM maps, (ii) the NMR-based identification of these structural elements along the sequence, in particular the residue-wise assignment of secondary structures, (iii) the unambiguous assignment of these sequence stretches to 3D structural features detected by EM, guided by NMR-derived distance restraints, and (iv) the joint refinement of the protein structure against NMR data and an electron-density map.

We show the feasibility of this approach to large protein assemblies with the example of the dodecameric 12 × 39 kDa (468 kDa) large aminopeptidase enzyme TET2 from *Pyrococcus horikoshii* which forms a large hollow lumen encapsulating twelve peptidase active sites.^15,16^ We use EM maps with resolutions of 4.1, 6 and 8 Å as well as secondary structure and distance information from MAS NMR and solution-NMR. Despite its large subunit size of 353 residues, larger than any protein assigned to date by MAS NMR, we achieved a near-complete NMR resonance assignment in a reliable manner, which allows the residue-wise determination of the backbone conformation, as well as hundreds of experimental distance restraints involving the protein backbone and side chain methyl groups, but by themselves these data are insufficient for structure determination. An automated approach that jointly exploits the electron density map, NMR secondary structure and distance restraints is presented, that allowed us to obtain the structure of TET2 at a backbone root-mean-square-deviation, RMSD, of 0.7 Å to the crystal structure.

Importantly, we show that the NMR/EM approach provides near-atomic-resolution structures even if only moderate-resolution EM data (up to 8 Å resolution) are available, an often-encountered situation that prohibits structure determination from EM data alone. Our structure provides insight into regions that had not been visible in previous crystal structures of TET2, in particular a conserved loop region in the catalytic chamber of this enzyme machinery.

## Results

### High-resolution NMR and cryo EM provide structural information on different length scales

Any investigation of structure, interactions or dynamics by NMR-based methods is bound to the availability of well-resolved spectra and the ability to assign the observed peaks to individual atoms along the protein sequence. Figure 1A and B show MAS NMR spectra of the protein backbone amides and solution-NMR of Ile and Val methyl groups in TET2, respectively. Despite the large number of residues per TET2 subunit, we obtained well-resolved spectra and high sensitivity. In solution, methyl spectra can be observed that are very similar to the corresponding MAS NMR spectra obtained. This shows that the sample states are structurally comparable, which allows combining the data from solution-NMR and MAS NMR in a straightforward manner (see Figure S1). To obtain atom-specific assignments of these spectra we collected a series of three- and four-dimensional ^13^C-detected and ^1^H-detected MAS NMR assignment spectra, which correlate the frequencies of neighboring atoms, thereby allowing to connect them along the protein backbone and out into the side chains (Figure S2A, B). Additional spectra recorded on three different samples with amino-acid-type specific ^13^C-labeling (either only LKP (Leu, Lys,Pro), GYFR or ILV residues) provided simplified spectra, which served as convenient starting points for manual assignment (see Figure S2C-F, Tables S2 to S4 and Methods for details). Together, these spectra were sufficient to obtain by manual analysis a near-complete assignments of TET2, ca. 85% of the backbone and 70% of the side chain heavy atoms (Figure S3). Residues for which no assignments were obtained are restricted to N- and C-termini and loops, a finding that can be attributed to the lower efficiency of the MAS NMR experiments for such dynamic parts. The assignment of 69 Ile, Leu, Val methyl groups was achieved through a combination of a mutagenesis-based strategy in solution, reported previously^17,18^, and the ^13^C frequencies obtained from MAS NMR assignment experiments. Even though TET2 is larger than any protein assigned to date to a similar extent by MAS NMR, the approach is sufficiently robust to obtain the near-complete assignment even in a fully automatic manner with the program FLYA^19^ (Figure S4), showing that reliable assignments can be obtained without time-consuming manual spectra analysis. The assigned chemical shifts allow the determination of *ϕ* and *ψ* backbone dihedral angles, and thus the secondary structure from a database approach, TALOS-N^20^. NMR-detected secondary structure elements are displayed in Figure 1D.

**Figure 1:**
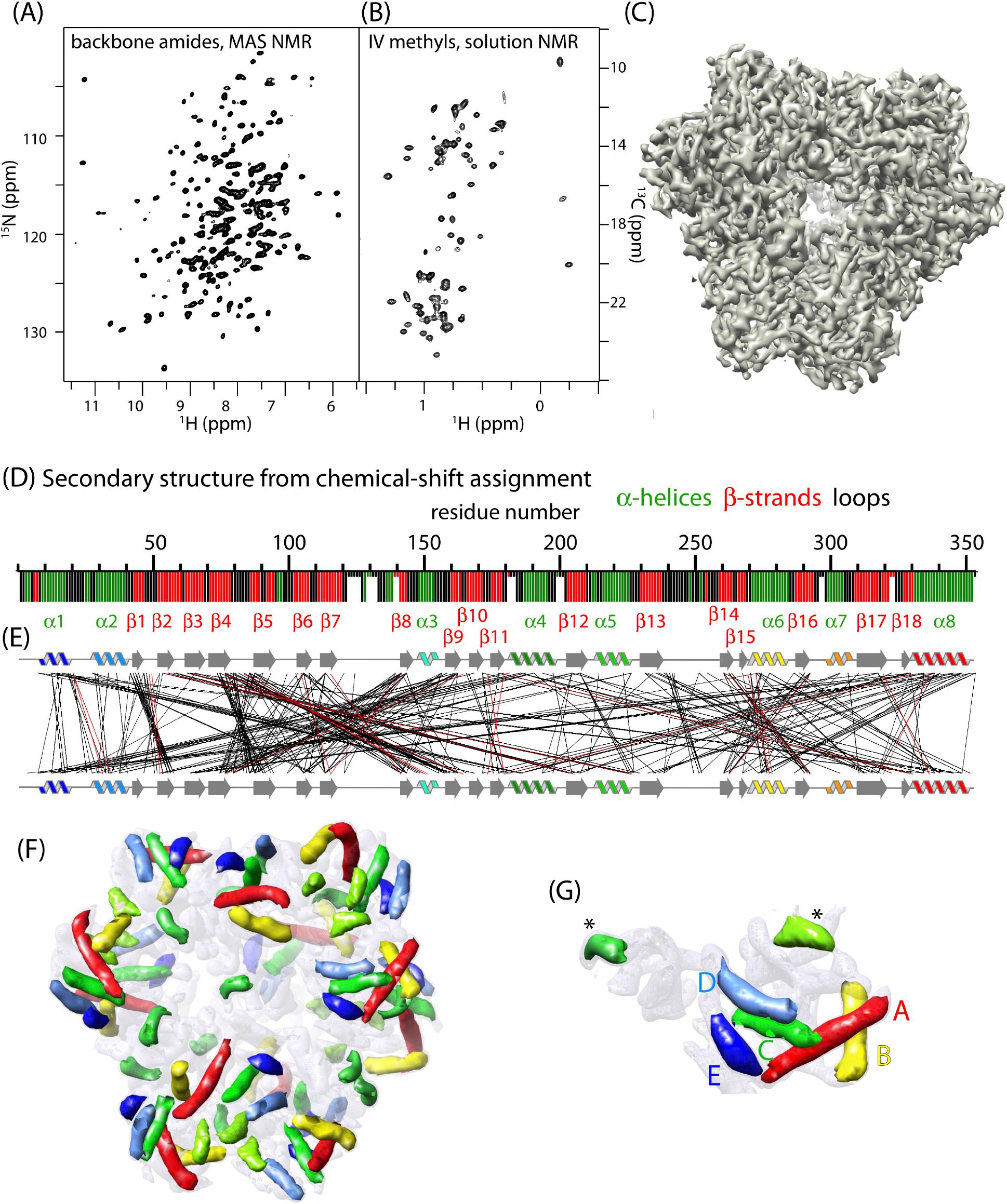
Experimental NMR and EM data of the 468 kDa TET2 assembly. (A) MAS NMR and (B) solution-NMR spectra of TET2, showing high resolution despite the large subunit size. (C) Experimental 4.1 Å resolution cryo-EM electron-density map. (D) Secondary structure of TET2, derived from MAS NMR resonance assignments and the TALOS-N software^20^, shown as a function of the residue number. Residues shown with shorter bars were not assigned, and the secondary structure assignment results from a database approach in TALOS-N. (E) Experimentally detected intra-subunit long-range distance restraints from solution-NMR and MAS NMR, displayed through lines connecting residues in close spatial proximity. Note that part of these distance restraints were spectrally ambiguous, i.e., could be assigned to several atom-atom pairs, and was rendered unambiguous throughout the structure calculation approach (displyed in red). See Table 1 for restraints statistics. All NMR experiments performed in this study and acquisition parameters are listed in Tables S2 to S5. (F) *α*-helices detected by the *helixhunter2* software^24,46^ in the EM map truncated to 8 A resolution. Symmetry-related *α*-helices are shown in equal colors. Additional *β*-sheet parts, automatically detected by *gorgon*^47^ are shown in Figure S9A. (G) Zoom on one subunit, identified by a clustering analysis (Figure S9). The five longest *α*-helices, used for the initial structure calculation steps are labeled with A to E in order of decreasing length (see Table S1).

In order to gain tertiary structure information, we measured short inter-atomic contacts through solution-NMR 3D nuclear Overhauser effect spectroscopy (NOESY) spectra and a 3D MAS NMR radio-frequency driven recoupling (RFDR) experiment (Figure S5). The solution-NMR data identify spatial proximity between Ile^*δ*1^ and Val^proS^ methyl groups; a second NOESY spectrum, recorded on a sample in which different subunits were labeled in different methyl groups, furthermore allowed filtering out inter-subunit methyl-methyl contacts (Figure S5E-G). The MAS NMR experiment provides distances between backbone amide sites and between amides and methyls of Ile, Val and Leu side chains, as well as contacts between such methyls in a single 3D MAS NMR experiment. ^13^C-^13^C distances were additionally measured in the selectively LKP-labeled sample. Figure 1E and Table 1 show all distance restraints identified by NMR, including 471 spectrally unambiguous ones, i.e., of which the cross-peak frequencies pointed unambiguously to one atom pair and 45 spectrally ambiguous ones, i.e., which could not be assigned to a single atom pair (but which could be unambiguously assigned along the structure calculation procedure, see below).

**Table 1.**
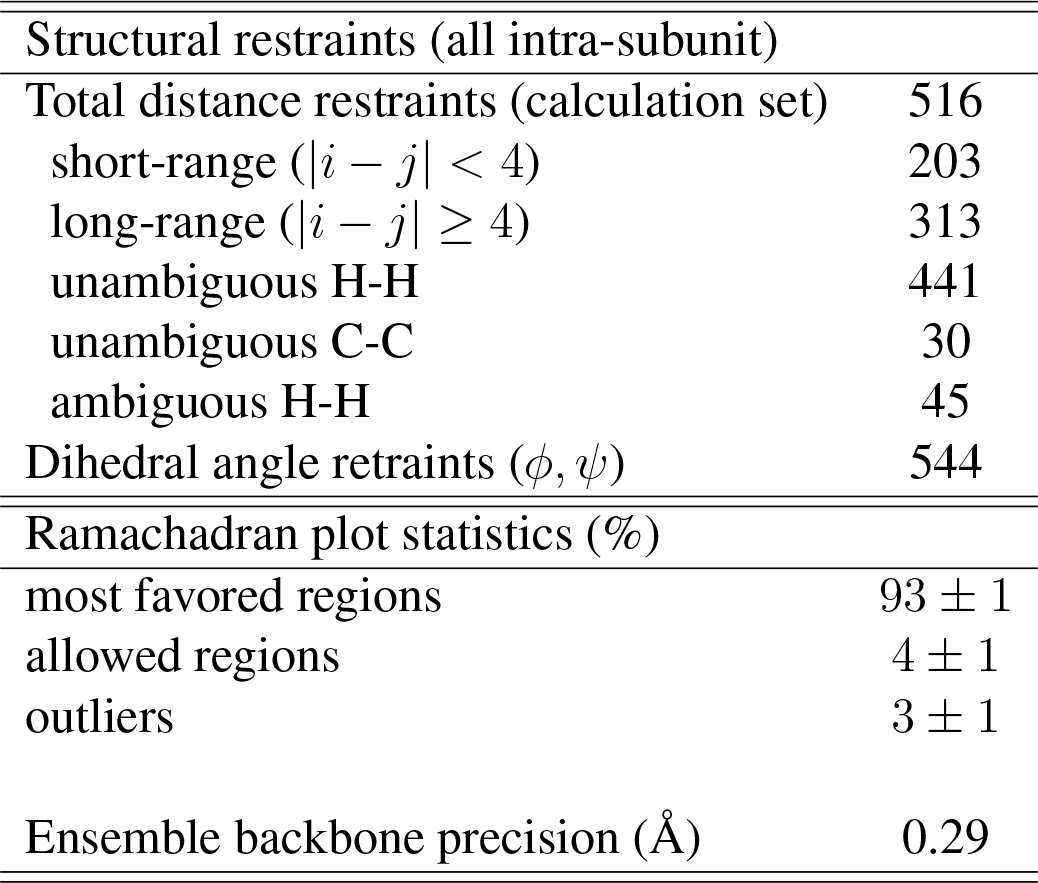
Number of distance and dihedral-angle restraints from NMR spectroscopy (top), and structure statistics of the final structure ensemble (20 structures) obtained from all NMR data and the 4.1 Å cryo-EM density map.

Attempts to determine the TET2 subunit structure from these NMR data using a standard structure-determination protocol based on restrained MD, CYANA^21^, failed in achieving convergence to the correct structure (backbone RMSD to the mean of the ten lowest-energy structures and to the crystal structure above 10 Å, see Figure S6). In these NMR-only structure calculations, many local structural elements, e.g., the local packing of two *α*-helices or *β*-strands, are formed, but the positions of these elements relative to each other in the tertiary fold remained poorly defined, thus calling for additional data.

Figure 1C shows the cryo-EM map of TET2 at 4.1 Å resolution, calculated from 27,130 single particles with the software RELION (see Methods section for details and Figure S7). We have attempted to build the structure of TET2 from this electron density map using *phenix.modelJojnap*^22^, in the crystallographic software suite Phenix.^23^ The program was able to trace the map, with better success when the symmetry of the particle was calculated from the cryo-EM map rather than inferred from the biological unit derived from the X-ray structure. However, the map quality was too low for phenix to assign the sequence and thus build a model (Figure S8). While it likely would have been possible to solve the structure at 4.1 Å by careful manual building, or by using other tools, the failure of this workhorse of crystallography model building illustrates the complexity of building a *de novo* model even at a resolution of 4.1 Å, which currently is among the high-resolution range for cryo-EM. This finding also highlights that, in all likelihood, succeeding with model building rapidly evaporates with decreasing resolution.

Despite the inability to trace the backbone chain in the density, a number of features, in particular *α*-helices, are readily recognized in the electron density, even at lower resolution (up to 8 Å) and in a fully automatic unsupervised manner ^24^. Figures 1F, G show the 12 × 7 *α*-helices that could be automatically detected from the electron density map of TET2 truncated to 8 Å resolution, and Figure S9A shows additionally detected *β*-sheet regions. NMR and EM thus provided structural information about TET2 on very different length scales, but neither of the two techniques by themselves allowed to determine the structure at the atomic scale, owing to the lack of long-range distance information in NMR, and the inability to build a robust model from the electron density.

### An automated approach for integrated EM/NMR structure calculation

We developed an approach for the combined use of EM and NMR data, keeping in mind that it should be applicable to cases where only medium-resolution EM maps (between 6 and 10 Å) would be available. Its central idea is the fact that certain structural elements can be identified even in such medium-resolution EM maps and by unsupervised automatic algorithms. The arrangement of these structural elements in 3D space, i.e., the knowledge of the positions of even a small subset of residues relative to each other, potentially brings very useful long-range structural information, and it is exactly such long-range information that would be of enormous benefit in NMR-based structure calculations.

Placing sequence stretches into the density can be viewed as a combinatorial assignment problem, in which sequence elements with known properties (e.g., *α*-helices, *β*-strands or large side chains that can be recognized in EM maps) are assigned to electron-density features with the matching shape. The most straightforward case is the one of *α*-helices, given the ease of detecting the *α*-helical electron-density features automatically, and assigning *α*-helical structures to sequence stretches from NMR data (Figure 1F,G and 1D, respectively).

Briefly, our approach, outlined in Figure 2, consists of three steps. In the first step, after detection of the *α*-helices, the various possibilities of placing *α*-helical sequence stretches into *α*-helix density features are systematically and combinatorially tested, performing with each of these helix-to-density assignments a restrained-MD type structure calculation with CYANA. This type of calculation follows standard procedures adopted in any NMR-based structure calculation, but the key difference is that in addition to NMR-derived distance restraints the *α*-helices are placed at precisely defined positions relative to each other. Subsequently, the correct helix-to-density assignment is identified through objective criteria that compare the resulting structural models to experimental NMR and EM observables. With the correct helix-to-density assignment at hand, steps 2 and 3 of our approach refine this model by iterative addition of restraints from the EM map as well as from NMR distance restraints that could be unambiguously assigned based on the initial structural models (see below).

**Figure 2:**
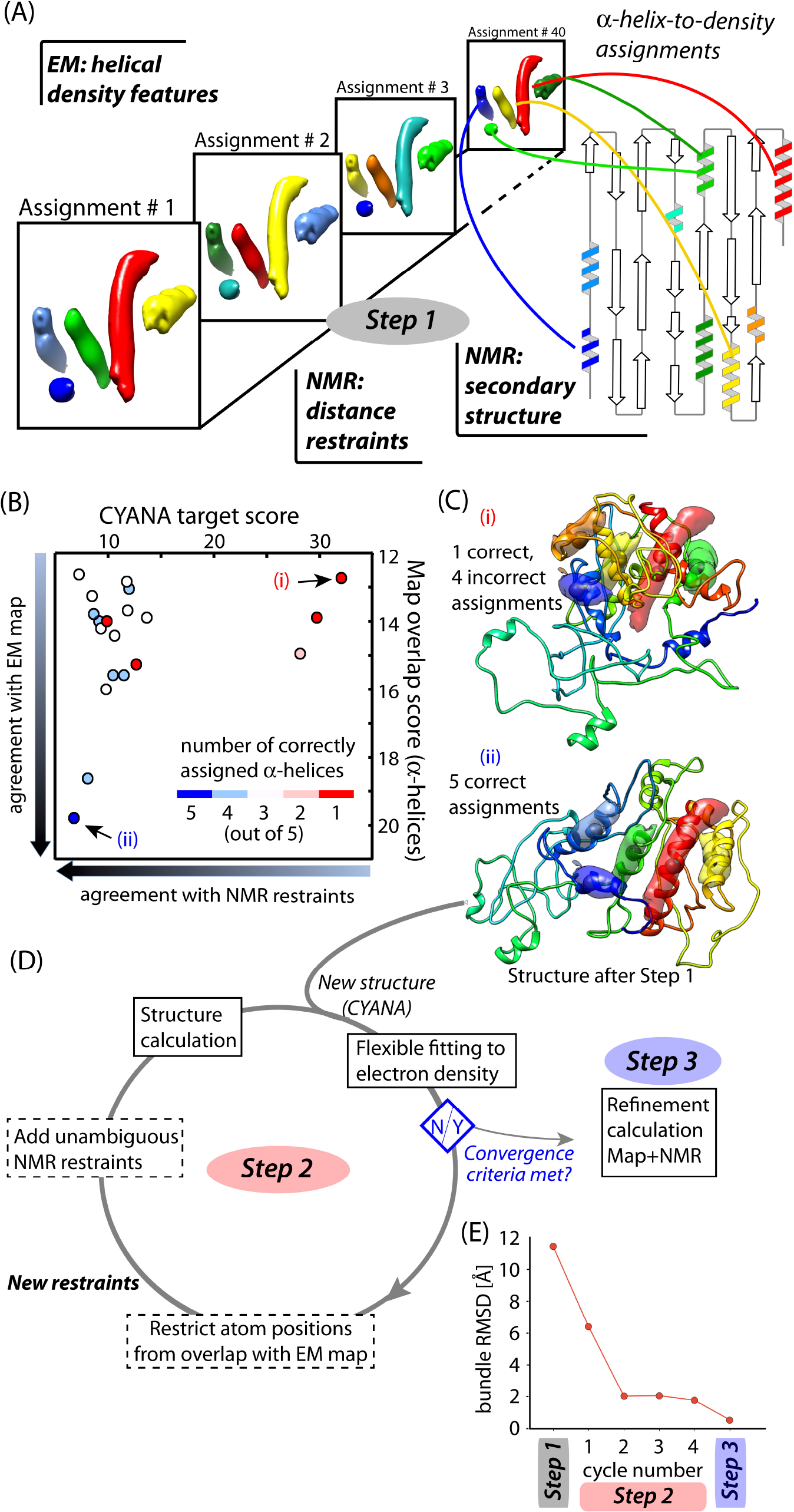
Integrated NMR/EM structure determination approach. (A) In step 1, forty different assignment possibilities of the five longest helical stretches in the sequence (TALOS) to cylindrical (helical) densities (”*α*-helix-to-density assignments”) are used in regular NMR-type structure calculations (CYANA). (B) Ranking of the twenty solutions from these structure calculations by the CYANA target score and the overlap of the *α*-helices with the cylindrical density features. Each point represents the average of the 10 lowest-energy structures. (C) Two cases are shown with incorrect (top) and correct (bottom) assignments, clearly showing that incorrect *α*-helix-to-density assignment is not compatible with good map overlap. For simplicity, only the lowest-energy structure is shown. The structure with the correct helix-to-density assignment (labeled with (ii)) has a backbone root-mean-square-deviation, RMSD, to the crystal structure of 7.4 Å, and a backbone bundle-RMSD computed from the 20 lowest-enerty models of 11. 4 Å. (D) In step 2, the structure of the TET2 monomeric subunit with the correct *α*-helix-to-density assignment is iteratively refined by flexible fitting into the EM map (truncated at 8 Å resolution), and CYANA calculations with an increasing number of unambiguous NMR restraints and restraints from the fit in the electron density. After convergence, defined by at least three cycles with an RMSD-difference below 10 % (shown in (E)), the structure is refined against the EM map of the entire dodecameric ensemble and all NMR restraints, using XPLOR-NIH, using maps with increasing resolution (8 Å, 6 Å, 4.1 Å; Step 3). (E) Root-mean-square-deviation (RMSD) of the structures at different steps of the protocol, relative to the mean structure.

An important consideration for the combinatorial procedure of step 1 is that the number of possible helix-to-density assignment combinations increases exponentially with the number of *α*-helices. Considering that the eight *α*-helices found in the sequence by NMR can be placed into the automatically detected helix features in the map (seven per subunit), including two orientations in each helix, there are 8! × 2^7^ ≈ 5 · 10^7^ possibilities (see Table S1 and discussion therein). Even more combinations are possible when considering that it is not straightforward to unambiguously identify which helical densities belong to the same subunit. Performing structural calculations with each of these assignments would be computationally very expensive.

A computationally manageable number of assignments can be achieved by taking into account some simple experimental observables. First, we used a cluster analysis to identify *α*-helical densities which are closest in space as belonging to the same asymmetric unit, and ascribed these five helical densities to the same subunit (see Figure 1G and Figure S9). Furthermore, we explicitly disregarded the polarity when assigning helical stretches to densities, by enforcing that only the central residue of a given *α*-helical sequence stretch is fixed to the center of an *α*-helical density, rather than the entire helix. Lastly, we discarded all those assignments for which the lengths of the *α*-helices along the sequence did not match the length of the helical density (see Table S1). With these assumptions and simplifications, only forty possible assignments remain and we performed forty CYANA structure calculations with these helix-to-density assignments (five residues restrained at fixed positions relative to each other), along with the backbone dihedral-angle and the spectrally unambiguous NMR distance restraints.

Identification of the correct helix-to-density assignment was based on two criteria, (i) the CYANA target function, which reflects the agreement of the structure with the distance and dihedral-angle restraints, and (ii) the overlap of the five *α*-helical stretches with the electron density. Incorrect helix-to-density assignments generally result in bent or twisted *α*-helices, and thus low overlap with the EM map and violation of NMR distances. Figure 2B displays these two scores for the twenty CYANA runs with the lowest CYANA score. A funnel-shaped distribution is observed for the various structures. Figure 2C displays two of these structures, in which either one or five *α*-helices were assigned correctly. The structure with the correct helix-to-density assignment scores best among all structures. While it has the topology approximately correct, its precision and accuracy are low, calling for further refinement steps.

The refinement stage of the protocol, an automated iterative procedure denoted here as Step 2 (Figure 2D), consists of iterative cycles of (i) a flexible fitting of the initial structure into the 8 Å EM density with iMODFIT^25^, and (ii) CYANA structure calculations. In each iteration, EM and NMR restraints were added: on the one hand, sequence stretches consistently located inside the 8 Å electron density were restrained to their positions, in the same manner as the helix-centers were initially fixed relative to each other (see Figure S10); on the other hand, the initial structural models allowed the unambiguous assignment of more NMR distance restraints, by excluding assignment possibilities corresponding to physically unrealistic long-range distances (see Methods). This iterative procedure was repeated until convergence of the bundle-RMSD (Figure 2E). In Step 3, a real-space structure refinement was applied in the XPLOR-NIH software^26,27^, using the full electron density map, first at 8 Å, then at 6 Å and lastly at 4.1 Å resolution, and all NMR restraints, calculating the full dodecamer. Figure 3 shows the evolution of the structure from the initial model to the final structure, which was obtained with high precision (bundle backbone RMSD over the 20 lowest-energy structures of 0.29 Å for one subunit).

**Figure 3:**
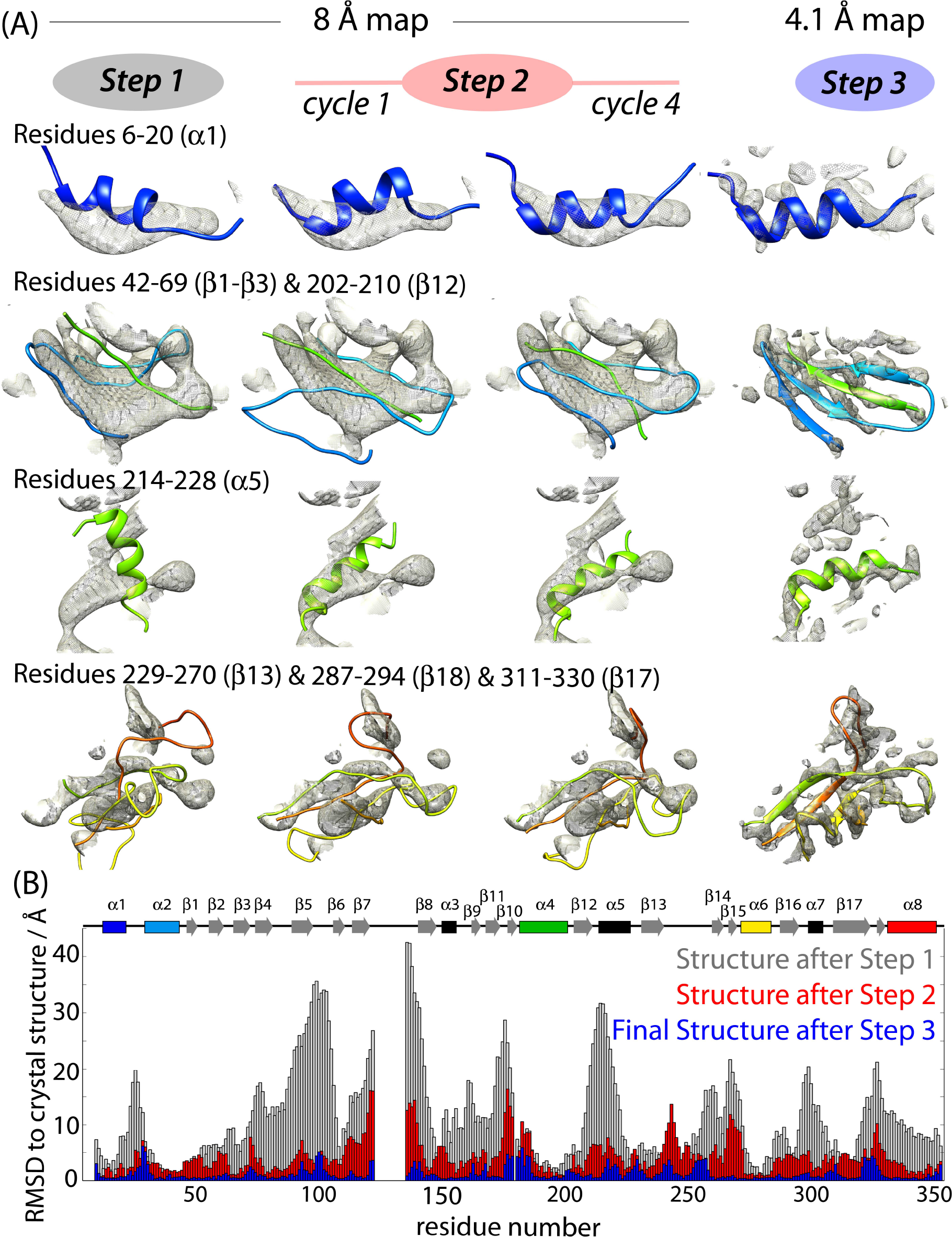
Structure refinement from NMR and EM data. (A) Zoom onto selected parts of the TET2 structure along the different steps of the structure determination process. The 3D EM map used at 8 Å resolution (steps 1 and 2) and 4.1 Å resolution (step 3) is shown as a gray surface. The structures shown correspond to the lowest-energy models generated by CYANA (Steps 1 and 2) or XPLOR-NIH (Step 3). A comprehensive view of the evolution of all structure elements is provided in Figure S11. (B) Residue-wise heavy-atom backbone RMSD relative to the crystal structure throughout different steps of the structure determination protocol.

The refined dodecameric structure (Figure 4) is in excellent agreement with the crystal structure. The dodecameric assembly forms a tetrahedral structure with ca. 12 Å wide openings, one on each of the four tetrahedron faces, providing access to the ca. 50 Å wide catalytic lumen. Our structure features a long, hithertho unobserved loop region in the catalytic chamber, comprising residues 119-138, shown in green in Figure 4C. Interestingly, even though the crystal structure has significantly higher resolution than the EM map (1.75 Å, ^16^ *vs*. 4.1 Å), this structural feature is much more clearly visible in the latter (Figure 4). Moreover, while this loop region appears relatively well defined at cryo-EM conditions, it is not observable at room temperature by MAS NMR, and we were unable to assign the resonances of the backbone of this loop (Figure S3E). These findings may suggest that the loop is dynamically disordered at room temperature, leading to low signal intensity in MAS NMR experiments, and to non interpretable density in crystal structures determined at cryogenic temperatures.

**Figure 4:**
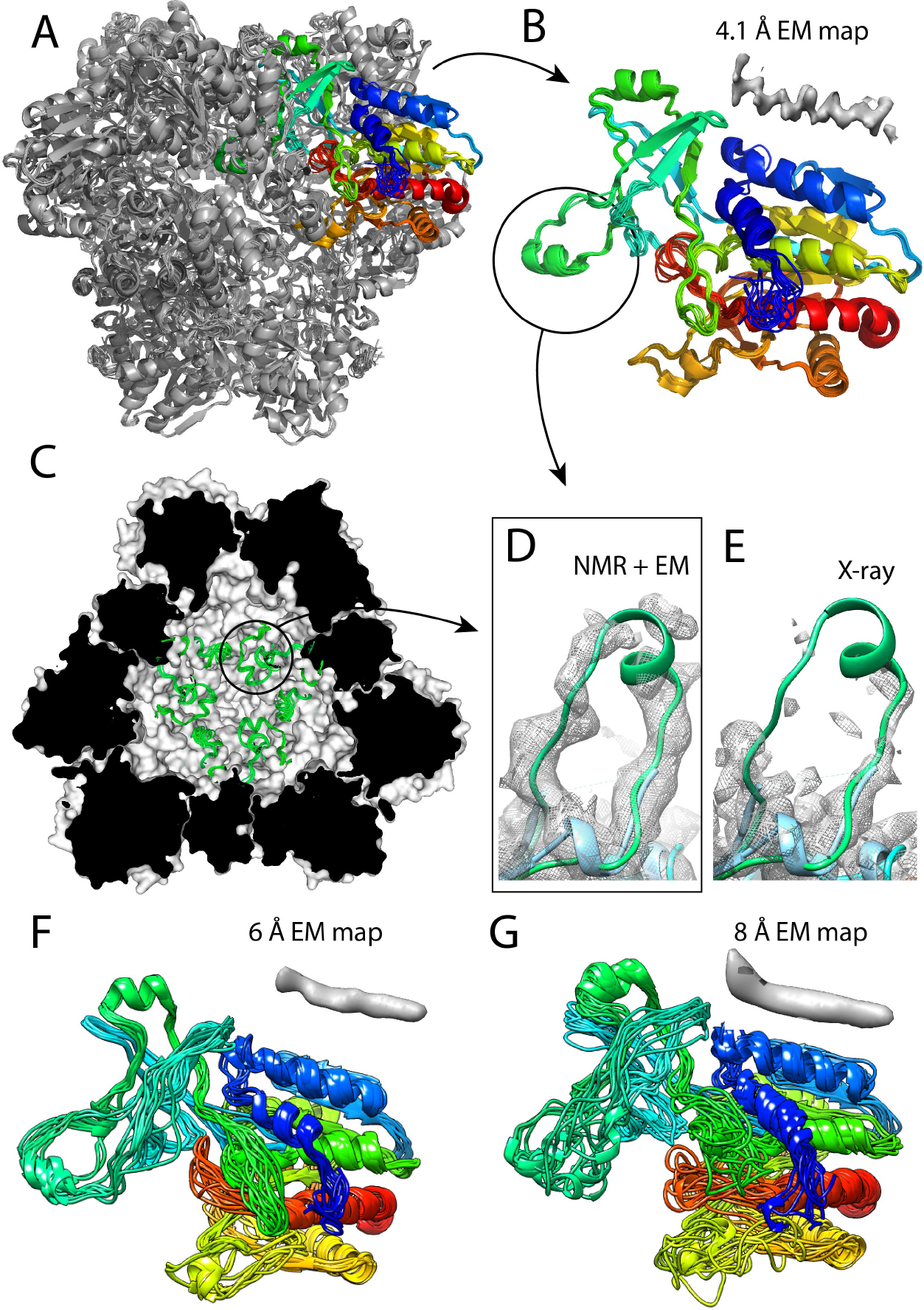
Structure of the TET2 dodecamer obtained from joint refinement of twelve subunits against NMR and EM data. (A) Overall view of the dodecamer, highlighting in color one out of the twelve modeled subunits. (B) Bundle of 10 structures of one monomer, with an accuracy of 0.7 Å (backbone RMSD relative to crystal structure). The loop region, residues 120-138, which has not been modeled in the crystal structure (PDB 1y0r) is encircled in (B), and shown in the view of the interior of the cavity (C) and as a zoom (D). The EM map around the loop region, seen by cryo-EM (D) is of significantly higher intensity than in the 100 K crystal structure (1y0r). The backbone structure in both panels is from the present NMR/EM study. (F, G) Results of NMR/EM structure calculation using only EM data truncated to 6 Å or 8 Å resolution, resulting in backbone RMSD to the crystal structure of 1.8 Åand 2.6 Å, respectively. The inserts above the structures in (B), (F) and (G) represent typical *α*-helical densities at the respective resolution levels.

An important consideration is how our approach performs with lower-resolution EM data. At 4.1 Å resolution, our EM map is better than the average of the maps deposited in recent years (average in 2018: 6.6 Å). We repeated our structure determination approach using maps truncated to either 6 Å or 8 Å in the refinement step. (Note that in Steps 1 and 2, our approach is anyhow based on a 8 Å map only.) Figures 4F and 4G show the refined structures with these two lower-resolution maps, which still have a decent accuracy, with backbone RMSD values to the crystal structure of 1.8 and 2.6 Å, respectively (calculated over Cα). We additionally explored a refinement with *phenix.real space-refine*^28^ to refine these structures furthermore, and obtained an even slightly higher structure accuracy (see Table S6 and Methods). These results highlight that the combined EM/NMR approach is able to provide near-atomic-resolution structures even in cases where the resolution is worse than the present-day average values, and for which *de novo* structure determination is currently far out of reach using EM methods only.

## Discussion

The recent “resolution revolution” in cryo-EM is progressively changing the way how the structures of biomolecular assemblies are determined, but currently *de novo* structure determination remains reserved to a minority of EM data sets. Even with the comparably favorable 4.1 Å resolution of our TET2 EM map we were unable to trace the polypeptide chain *de novo* using currently available tools (see Figure S8). The integration of complementary data from different techniques holds great promise for obtaining atomic-resolution structures even when any single method provides too sparse data. The present work is among the few examples of such an integration of EM and NMR data. Compared to these previous structures determined from NMR/EM data^12–14^, TET2 has a more than four times larger monomer size, being larger than any protein for which near-complete MAS NMR resonance assignments have been reported so far.

Our approach uses features that can be identified in a straightforward manner even in medium (8-10 Å) resolution EM maps, and in an automatic unsupervised manner, most commonly *α*-helices. Although not exploited here, also some *β*-sheet regions and bulky side chains could be readily identified in the EM maps (cf. Figure S9A) and exploited similarly as the *α*-helices used here. The inclusion of only very few such real-space restraints—in our case only five points along the sequence—turns out to be extremely powerful in driving the structure calculation towards the correct fold. Specific isotope-labeling schemes, in particular deuteration and methyl-labeling, were crucial as they allowed the unambiguous assignment of hundreds of NMR-distance restraints between residues distant in primary sequence, thereby enabling to use the NMR-score to identify the correct helix-to-density assignment, which is the first step to guide the structure towards the correct topology. A remarkable result of this study is the finding that even an EM map at 8 Å resolution allowed us to determine the fold accurately, whereas the *de novo* modeling even with a 4.1 Å resolution map failed.

Data from other sources can be readily included into this integrated approach. The distance restraints obtained here from solution-state and MAS NMR may be complemented or replaced by distance information obtained from co-evolution data^29–31^, cross-linking experiments,^32^ Förster resonance energy transfer,^33^ DEER electron paramagnetic resonance^34–36^, or NMR-detected paramagnetic relaxation enhancement (PRE) data.^37^ The secondary structure information obtained here from NMR chemical-shift assignments may be complemented with prediction algorithms in case of missing NMR assignments. Integrating such diverse data in structure determination will likely play an important role in structural studies of challenging biological assemblies forming oligomeric structures, such as capsids, tubes/needles/fibers, i.e., structures that generally are very difficult to study by crystallography.

## Methods

### 1 Sample preparation

#### Expression and isotope-labeling of TET2

TET2 from *Pyrococcus horikoskii* (UniProt entry O59196) was obtained by overexpression in *E. coli* BL21(DE3) cells with suitable isotope-labeled M9 minima, supplemented with ^15^NH_4_Cl in all cases as sole nitrogen source. Depending on the desired labeling scheme (outlined below), suitably labeled D-glucose (uniformly ^13^C_6_-labeled or 1,2,3,4,5,6,6-^2^H_7_,^13^C_6_-labeled or 1,2,3,4,5,6,6-^2^H_7_-labeled or unlabeled) was used as carbon source, and for the case of samples with amino-acid specific labeling, isotope-labeled amino acids or ketoacid precursors were added before induction with isopropyl *β*-D-1-thiogalactopyranoside (IPTG, 1 mM in final culture).

For MAS NMR, six different samples were used in this study with different labeling. Sample 1: u-[^13^C,^15^N]. Sample 2: u-[^15^N]-Gly,Tyr,Phe,Arg-[^13^C]. Sample 3: u-[^15^N]-Ile,Leu,Val-[^13^C]. Sample 4: u-[^15^N]-Leu,Lys,Pro-[^13^C]. Sample 5: u-[^2^H,^13^C,^15^N]. Sample 6: u-[^2^H,^15^N], Ile^*δ*1^,Leu^*δ*2^,Val^*γ*2^-[^13^CHD_2_], i.e., ^13^CHD_2_ methyl groups at Ile-*δ*^1^, Val-*γ*^proS^ and Leu-*δ*^proS^ positions in an otherwise perdeuterated background (the pro-S methyl is the one denoted as *γ*2 in valine and as *δ*2 in leucine).

For sample 1, ^13^C_6_-labeled glucose was used. For samples 2-4, unlabeled D-glucose was used. For sample 3 ^13^C_4_-2-ketobutyrate (60 mg/L of culture) was added 20 minutes prior to induction and ^13^C_5_-acetolactate (300 mg/L) was added 1 h prior to induction. The resulting Ile residues have ^13^C labels at the C”, C^*α*^, C^*γ*1^ and C^*δ*1^ sites; the leucines bear ^13^C at the *β*, *γ* and *δ* sites; the valines are uniformly ^13^C labeled^38^. For samples 2 and 4, the u-[^13^C,^15^N]-labeled amino acids were added 1 h prior to induction.^39^ For sample 5, D_2_O-based M9 medium was used, with ^2^H_7_,^13^C_6_-labeled glucose. For sample 6, Ile^*δ*1^,Leu^*δ*2^,Val^*γ*2^ sites were labeled by addition of 2-ketobutyrate (60 mg/L; 20 min prior to induction) and acetolactate (200 mg/L; 1 h prior to induction), with ^13^CHD_2_-groups at the relevant positions^38,40^. For the production of the deuterated, methyl-labeled samples, cells were progressively adapted, in three stages, over 24 h, to M9/D2O media containing 1 g/L ^15^ND_4_Cl and 2 g/L ^2^H_7_ glucose (Isotec). In the final culture, the bacteria were grown at 37°C in M9 media prepared with 99.85% D_2_O (Eurisotop).

Samples 1–4 were used for resonance assignment with ^13^C-detected MAS NMR experiments (see Table S2). Sample 5 was used for ^1^ H-detected MAS NMR assignment experiments, listed in Table S4. Sample 6 was used to collect ^1^H-^1^H distance restraints between methyl and amide groups by RFDR MAS NMR (see Table S5). Sample 4 was furthermore used to collect ^13^C-^13^C distance restraints.

For solution-state NMR NOESY experiments, a uniformly deuterated sample with ^13^CH_3_ groups at Ile-*δ*^1^ and Val-*γ*^proS^ sites was used (sample 7). We refer to this labeling as u-[^2^H], u-[^15^N], Ile^*δ*1^,Val^*γ*2^-[^13^CH_3_]). This sample was used for collecting methyl-NOESY spectra, listed in Table S5. Three additional samples were used for the identification of intermolecular contacts, labeled either with u-[^2^H,^15^N],Ile^*δ*1^,Thr^*γ*2^,Val^proS^-[^13^CH_3_] (sample 8), or u-[^2^H,^15^N],Met^*ϵ*^,Ala^*β*^,Val^proR^-[^13^CH_3_] (sample 9), and a sample in which these two types of subunits were re-assembled (sample 10), as described in Figure S5E-G. In the present work, the detection of inter-subunit contacts, however, turned out to be not critical, and these three samples and the according NOESY data sets were used only to exclude eight inter-subunit contacts.

#### Purification of TET2

Cells were harvested by centrifugation and broken in a microfluidizer (15 kpsi, 3 passes) in 50 mM Tris-HCl/150 mM NaCl buffer. The total protein extract from disrupted cells was heated to 85°C for 15 minutes and the lysates cleared by centrifugation (17000 g, 1 h, 4°C). The supernatant was loaded onto a 6-ml Resource Q column (GE Healthcare) equilibrated in 100 mM NaCl, 20 mM Tris-HCl, pH 7.5. After washing with 3 column volumes, the protein was eluted with a linear salt gradient (0.1 - 0.35 M NaCl in 20 mM Tris-HCl, pH 7.5), followed by a size exclusion chromatography step (HiLoad 16/60 Superdex 200, GE Healthcare). Sample purity was verified by SDS-PAGE, and protein-containing fractions were pooled and concentrated.

#### NMR samples

Samples used for the solution-state NMR NOESY experiment were at a protein subunit concentration of 1 mM (i.e., a concentration of dodecameric particles of 83 *μ*M) in a D_2_O-based buffer (20 mM Tris, 20 mM NaCl, pH 7.4) in a 5 mm Shigemi tube.

Samples for MAS NMR were prepared by concentrating TET2 to 10 mg/mL in H_2_O-based buffer (20 mM Tris, 20 mM NaCl, pH 7.4) and then adding 2-Methyl-2,4-pentanediol (MPD), a commonly used crystallization agent, in a 1:1 (v/v) ratio. A white precipitate (possibly of nanocrystalline nature) appeared immediately after mixing of the solutions. The samples were filled into either 1.3 mm (Bruker), 1.6 mm (Agilent), 3.2 mm Agilent or 3.2 mm (Bruker) rotors, using an ultracentrifuge filling device adapted for a Beckman SW32 ultracentrifuge. Typically, samples were filled for ca. 1 h at ca. 50000 x g. Rotors were glued with two-component epoxy glue to avoid loss of solvent.

#### Sample for cryo-electron microscopy

A sample of u-[^2^H,^13^C,^15^N]-labeled TET2 was used for cryo-electron microscopy. The isotope labeling, although unnecessary for cryo-EM, was useful to record ssNMR spectra with the same sample, in parallel to cryo-EM as a sample quality control. Three and a half microliters of sample were applied to 2:1 glow discharged Quantifoil holey carbon grids (Quantifoil Micro Tools GmbH, Germany) and the grids were frozen in liquid ethane with a Vitrobot Mark IV (FEI, the Netherlands).

### 2 NMR spectroscopy

All solution-NMR experiments were performed at a temperature of 50°C, either with a Varian INOVA 800 MHz spectrometer (for the H-H-C NOESY) or a 950 MHz Bruker Avance 3 spectrometer, both equipped with cryogenically cooled probes. MAS NMR experiments were performed at an effective sample temperature of ca. 28^°^C, measured from the bulk water resonance and the resonance of MPD at 4.1 ppm. MAS NMR assignment experiments were recorded either on a Agilent 600 MHz VNMRS spectrometer equipped with a triple-resonance 3.2 mm probe (for ^13^C-detected experiments) or a 1.6 mm Fast-MAS probe (for all reported ^1^H-detected experiments at a MAS frequency below 40 kHz), or a Bruker 1000 MHz Avance spectrometer equipped with a 3.2 mm probe (only one 3D experiment, see Table S2. All additional ^1^H-detected experiments at MAS frequencies >50 kHz were recorded with a Bruker 600 MHz Avance 3 spectrometer equipped with a 1.3 mm MAS probe tuned to ^1^H, ^2^H, ^13^C and ^15^N frequencies. Acquisition parameters of all experiments are reported in Tables S2 to S5.

#### NMR resonance assignment

Assignment of NMR resonances by MAS NMR was performed with a suite of 3D and 4D experiments with either ^13^C detection or ^1^H detection, listed in Tables S2, S3 and S4.

Inspection of the spectra and manual peak picking was performed with the CCPNMR software^41^. Near-complete resonance assignments were achieved by manual analysis. Hereby, the two 4D spectra, CONCACB and CANCOCX, were crucial to unambiguously identify neighboring spin systems. They share three frequencies (CA, N, CO), and the combination of the two spectra thus allows unambiguously connecting five or six frequencies. Comparing patterns in these spectra, as well as in 3D NCACX, NCOCX and NCACB spectra allowed the sequential connection of such 5-tuples of frequencies. Assignments of the side chains were obtained from the threedimensional NCACX, NCOCX and CCC experiments. The assignment of all reported heavy atoms was achieved from ^13^C-detected experiments only. The additional ^1^H-detected experiments were used to assign the amide-^1^H frequency.

Following the manual assignment we investigated the ability to obtain fully automatic assignments. Peaks in all spectra listed in Tables S2–S4 were picked manually (numbers of picked peaks are listed in Tables S2 to S4), and used as input for the automated assignment algorithm FLYA^19^, implemented in CYANA version 3.97^21^. The output of the automatic assignment procedure is shown in Figure S4.

This suite of experiments allowed the assignment of the heavy atoms (^13^C, ^15^N) and amide ^1^H. The additional assignment of methyl groups was achieved by a mutagenesis-based solution-NMR assignment approach, reported previously for Ile ^17^ and Val ^18^. These assignments were confirmed and extended to Leu by the ^13^C assignments obtained by MAS NMR.

In cases where several methyl groups have the same ^13^C frequency, this information was insufficient by itself. Thus, we furthermore used the information from the RFDR MAS NMR experiment, in particular cross-peaks from the methyl group to the amide backbone site. All resonance assignments are reported in the BioMagResBank (BMRB ID 27211).

#### NMR distance restraints measurement

Table S5 summarizes all acquisition parameters of solution-state and MAS NMR experiments used for determining distance restraints. Two types of solution-state NMR NOESY spectra were collected. In a first approach, a u-[^2^H, ^15^N], Ile-[^13^CH_3_]^*δ*1^, Val-[^13^CH_3_]^pro-S^ labeled sample was used to collected NOESY distance retraints with a 3D H-H-C edited experiment.^18^ In order to exclude inter-subunit contacts, which might induce errors in single subunit structure calculations, we additionally measured two 3D H-C-C NOESY experiments: one on a sample in which different subunits were labeled differently (sample 10; see Figure S5E) and two samples where all subunits were labeled identically (sample 8). Eight inter-subunit contacts detected in this experiment were excluded from the calculation.

MAS NMR distance restraints were obtained through a 3D ^1^H-^15^N/^13^C-^15^N/^13^C time-shared RFDR experiment and a 2D ^13^C-^13^DARR experiment (see Table S5 and Figure S5).

### 3 Cryo-electron microscopy

The sample was observed with a FEI Polara at 300 kV. Movies were recorded manually on a K2 summit direct detector (Gatan Inc., USA) in super resolution counting mode at a nominal magnification of 20,000 (0.97 Å/pixel at the camera level) for a total exposure time of 4 s and 100 ms per frame resulting in movies of 40 frames with a total dose of ~40 e-/Å^2^. 90 movies were manually recorded with Digital Micrograph (Gatan Inc., USA). Movies were first motion corrected with Digital Micrograph (Gatan Inc., USA) resulting in micrographs like the one on Figure S7. 1643 particles were picked semi-automatically with boxer from the EMAN suite^42^ and 2D classified in 16 classes in RELION 1.4^43^. The best-defined 2D class averages were used to autopick particles in all the micrographs with RELION resulting in a total of 30407 particles. CTF estimation with CTFFIND3^44^, particle extraction in boxes of 200 × 200 pixels and preprocessing were done in RELION as well. The data set was first cleaned by 2D classification leaving 27130 particles. From the best 2D classes, a low resolution *ab initio* 3D model was generated using the RIco server^45^ imposing tetrahedral symmetry. The latter was then used as a starting model to refine all particle orientations in RELION (with tetrahedral symmetry imposed). Further 3D classifications were attempted but did not result in any improvement of the structure. The final 3D reconstruction has a resolution of 4.1 Å at FSC = 0.143.

### 4 Structure calculation

The protocol described in this work employed standard programs routinely used in NMR structure determination and EM fitting, namely CYANA (version 3.97)^21^, XPLOR-NIH (version 2.44.8),^26^ iModFit (version 1.44),^25^, helixhunter2 ^24,46^, Phenix as well as in-house written scripts in python language (available from the authors upon request).

#### Step 1a: *α*-helix detection and assignment

The automatic detection of the *α*-helix and *β*-sheet regions from the cryo-EM map was done with the 8 Å resolution map using helixhunter2 ^24,46^ and *gorgon*.^47^ *helixhunter1*, was able to detect only the five longer alpha helices, while both *helixhunter2* and *gorgon* added two shorter *α*-helices and the latter also detected *β*-sheet regions that are shown in Figure S9A. We used only the five longest *α*-helices (and their eleven symmetry mates across the dodecamer), denoted A-E in Figure 1G and Table S1, for generating structure restraints used in Step 1 of the protocol.

While not strictly necessary in this protocol, for reasons of computational efficiency it is helpful to be able to identify which *α*-helices belong to a given TET2 monomer, as it allows reducing the number of helix-to-density assignments if only helical densities within one subunit need to be considered. We clustered these 12 × 5 helical densities into groups corresponding to individual subunits by an automatic approach based on the “density-based spatial clustering of application with noise (DBSCAN)” algorithm that was implemented in the Sklearn python package (https://pypi.org/project/sklearn/#history), and the outcome is shown in Figure S9. In essence, the underlying assumption is that the *α*-helices that are closest to each other belong to the same subunit. For the case of the five helices we considered in TET2, this assumption turns out to be correct. Note, however, that our approach does not rely on the validity of the assumption that the closest helices belong to one subunit; any other combinations, including (erroneous) assignments of one chain to densities belonging to different subunits can equally be considered in the CYANA calculations. We have tested a number of such erroneous helix-to-density assignments, and all these calculations yielded very high CYANA scores and could be discarded on this basis (data not shown). Thus, erroneous assignment of one chain to different subunits is detected through lack of convergence along the protocol.

Table S1 shows the helix-to-density assignment possibilities which we have taken into account in Step 1, based on the lengths of the *α*-helices. In addition to considering the helix lengths, all the physically meaningless assignments in which the same a-helix would be placed at two or more locations at the same time are filtered out. The forty possible assignments, listed in Table S1) were used to generate lists of residue numbers (the centers of the NMR-detected *α*-helixces) assigned to coordinates in space (the centers of the five *α*-helical densities). These inter-helix distance restraints were used in the subsequent CYANA calculations.

#### Step 1b: CYANA structure calculation and ranking assignments

Forty CYANA calculations (version 3.97)^21^ were performed, using as restraints the unambiguous distance restraints, backbone dihedral-angle restraints and the restraints of helix-centers to helical-density centers. In all CYANA calculations performed in this study, only one chain (353 residues) was used, and the dodecamer was built from twelve chains only in Step 3. Because CYANA works in dihedral-angle space in a protein-internal frame, rather than in real space, the implementation of the real-space helix restraints required translating the absolute-space information into relative-space information. In practice, distance restraints were applied between all atoms that were to be fixed in real space, e.g., between the five helix-centers in Step 1. For practical purposes, this method results in fixing the relative orientation and distance of these points, but leaves translational and rotational freedom to the whole protein, which is inconsequential for the resulting structure.

For each helix-to-density assignment, one-thousand conformers were computed using the standard simulating annealing protocol in CYANA with 4000 torsion-angle dynamics steps per conformer. Typical computation times for generating these 1000 conformations was ca. 8.3 minutes on an Intel 8-core desktop computer. Finally, the 10 conformers with the lowest final target energy function were retained, and their energy was used as one criterion for ranking of the assignments. This CYANA score is plotted as one of the axes in Figure 2B. Note that out of the forty CYANA runs, twenty were stopped due to divergent target function energy, showing that these helix-to-density assignments were incompatible with the other NMR data. The twenty remaining calculations are shown in Figure 2B.

The second criterion for selecting the correct assignment was the goodness of the fit between the electron density map and the structure, evaluated only for the five helices considered in a given helix-to-density assignment. Basically, the idea is to evaluate whether all residues of the helix reside within the experimental electron density; incorrect helix-to-density assignment would lead to a distorted structure and therefore the helices would not fit in the electron density.

To implement this criterion in practice, electron-density maps were generated in silico from the lowest-energy CYANA model, using the backbone N, C*α*, C and O atoms of each of the residues in the five helices which were used in the given helix-to-density assignment (as identified by TALOS-N), using the module *molmap* in UCSF Chimera.^48^ We compared these in silico maps to an experimental map, which comprised only the five *α*-helices considered (colored and labeled A-E in Figure 1G, map truncated to 8 Å resolution). The module *measure correlation* in UCSF chimera was used to compute the correlation between the in silico maps (of each residue) and this experimental “helix-density” map. This correlation coefficient, obtained residue by residue, reflects the goodness of fit to the map. Examples are shown in Figure S10. As a global measure, we computed the percentage of residues that had a good overlap (> 0.7, in which 1.0 and 0.0 means good and bad overlap between the maps, respectively). This percentage is shown in Figure 2B as vertical axis.

#### Step 2: Iterative refinement

In this step, we performed iterative rounds of flexible fitting using a common tool used in EM (iModFit^25^) and an NMR-type structure calculation with CYANA. For the latter we used the information about the match to the EM map, similarly as the five helix centers were restrainted in Step 1. This procedure is described as follows.

The structure resulting from Step 1 was taken as a seed to perform a normal-mode based flexible fitting into the density map with the software iModFit v1.44.^25^ After this procedure, we identified those protein regions which resided well within the experimental electron density, in order to be able to fix those residues in the following CYANA calculations. This identification how well the residues correspond to experimental electron density was done in a similar manner as the goodness-of-fit criterion described in the preceding paragraph, using UCSF chimera modules *molmap* (allowing to generate density maps from a PDB file) and *measure correlation* (allowing the computation with an experimental map), as follows. We created an in silico map for each residue (backbone N, Ca, C, O only) and computed its correlation to the experimental EM map, truncated to 8 Å resolution, using the full map (rather than only the helices used above). Figure S10 shows this overlap score for each residue, as well as graphical examples. Residues with a good overlap score to the map (arbitrarily set as > 0.7), were restrained in the following CYANA structure calculation. To fix the atoms in real space using CYANA, even though CYANA operates in internal dihedral-angle space, the same procedure was applied as in Step 1, i.e., the atoms were constrained through distance restraints to all those other atoms that were also fixed in real space (using a tolerance of ±0.5*Å*). *Note that in practice it is of no importance where in absolute space the structure is placed, so long as the E atomic distances was generated using an in–house written python script, and was used in the same manner as NMR distance restraints,*

NMR distance restraints with ambiguous assignments, i.e., resulting from cross-peaks with frequencies that could be assigned to more than one atom-pair, can be disambiguated with the help of the intermediate structure obtained after the flexible fitting. Briefly, for cases with multiple possible assignments (based on NMR chemical-shift positions) we measured the corresponding distances in the intermediate structures for all those atom pairs that were possible assignment candidates based on their frequency. In cases where out of the possible candidates only one atom pair had short a (< 8 Å) distance, this assignment was retained in the subsequent CYANA calculation. Note that one may also use the restraints as ambiguous restraints in CYANA, but we found better convergence with this approach.

Furthermore, we also considered that some of the NOESY and RFDR distance restraints may be due to inter-subunit contacts. Introducing these restraints in a calculation that only considers one chain would necessarily lead to distance violations. To identify such cases (of which we had only 8 altogether), 3D HMQC-NOESY-HMQC spectra were recorded on a sample in which differently labeled subunit types were mixed in a 1:1 ratio and which allows recognizing inter-subunit cross-peaks (see Figure S5G-I) and compared to a uniformly labeled sample (see Figure S5E-G for details). Cross-peaks identified in this way as stemming from inter-subunit contacts were excluded from further analysis.

CYANA calculations in Step 2 used this growing list of restraints from the match to the map as well as the increasing list of unambiguous NMR distances along with the NMR dihedral-angle restraints. Only one subunit was used in these calculations; the explicit dodecamer was considered only in Step 3.

#### Step 3: Final real-space refinement

In the final refinement step, all dihedral-angle and distance restraints from NMR as well as the full EM map (in real space), at a resolution of either 4.1, 6 Å (Figure 4F) or 8 Å (Figure 4G), were used in a joint calculation of the full dodecamer using XPLOR-NIH ^26^ version 2.44.8. Xplor-NIH’s strict symmetry facility ^49^ was used to generate subunit coordinates with tetrahedral symmetry from those of a protomer subunit. Initially, protomer coordinates from Step 2 were moved as a rigid body to fit the full construct into the EM map, also allowing overall center of mass motion so that the dodecamer could be centered on the map. This starting structure was refined, allowing all protomer degrees of freedom under all restraints using the standard purely repulsive nonbonded energy term. In each refinement calculation 100 structures were calculated differing in random velocities given at the beginning of molecular dynamics simulated annealing.

The spatial restraining effects of the EM map were introduced by performing calculations against progressively higher resolution maps: the 8 Å map, followed by the 6 Å map, and finally the highest-resolution (4.1 Å) map. The fit structure was used as initial coordinates for the first calculation, and the lowest energy calculated structure from each step was utilized as the initial structure for the subsequent calculation. Finally, the outcome of the calculation using the 4.1 Å map was refined in implicit solvent using the effective energy function for Xplor-NIH (EEFx)^50^. For generating the structures shown in Figure 4F and G, this gradual refinement process was stopped before adding the higher-resolution maps (i.e., before adding the 6 or 4.1 Å resolution maps, respectively), in order to evaluate the quality of the structures resolution from lower-resolution EM data.

In a final stage, to locally refine the structure of the Zn_2_ active site, we introduced distance restraints to the zinc ions. The identity of these chelating side chains can be readily predicted from the sequence.^51^ At this stage of the protocol, the chelating residues (H68, D182, E213, E235, H323) were already close in space to each other, allowing the inclusion of explicit restraints between these atoms to model the zinc site using generous distances based on those found in crystal structures. Also in this region, restraints were added from residue 323 to residues 92 and 93 based on a subsequent reanalysis of unassigned RFDR cross peaks. The final XPLOR-NIH calculation was performed in implicit solvent with the 4.1 Å resolution map, all NMR restraints and the Zn-site restraints. For the calculation with the lower-resolution maps (6 Å, 8 Å), this refinement in implicit solvent has been omitted. The structure deposited as 6f3k in the PDB corresponds to the result of this refinement.

In addition, we also investigated whether the result can be further improved with phenix, a commonly used program in crystallography. The lowest-energy structures from the XPLOR-NIH runs, employing EM data at different resolution, were refined against the cryo-EM maps using *phenix.realspace-refine*^28^. We additionally used as restraints all distance measurements and secondary structure information from NMR. Different refinement schemes were tested, which all included five iterative cycles of rigid-body fit of individual chains and minimization of atomic positions, but also simulated annealing or morphing or local grid search. The results of these refinements are summarized in Table S6. We found simulated annealing to be superior to the two other approaches, as well as to not using it. Models obtained by performing Step 3 at 6 and 8 Å, respectively, were refined using the same approach, i.e. by five iterative cycles of real-space refinement including rigid-body fit of individual chains, minimization of atomic positions, and simulated annealing. Refinements were carried out both at 6 or 8 Å, and at 4.1 Å, to allow a fair comparison of the low resolution models with that determined by performing Step 3 at 4.1 Å. Correlation coefficients between the cryo-EM map and the model were calculated using *phenix.realspace-refine*, and correspond to the CCmask described by Jiang & Brunger^52^ based on map values inside a mask calculated around the macromolecule. Root-mean-square deviations between the various models and the biological unit derived from the X-ray structure (1y0r) were calculated using the *super* function in PyMOL.

### De novo structure building from the EM map with Phenix

We attempted to rebuild the TET2 structure *de novo*, by use of the 4.1 Å cryo-EM map only, using *phenix.modeLto-map*^22^, in the Phenix suite of crystallographic software ^23^. The program was able to trace the map, with better success when the symmetry of the particle was calculated from the cryo-EM map rather than inferred from the biological unit derived from the X-ray structure. However, the map quality was too low for phenix to assign the sequence and thus build a model. We tested a variety of building schemes, the results of some of which are shown in Figure S8).

### Data deposition

The NMR assignment has been deposited in the BioMagResBank (entry 27211). The 4.1 Å EM map has been deposited in the EMDB (entry 4179). The atomic model resulting from the three steps described herein, using in the final step the 4.1 Å EM map, has been deposited in the Protein Data Bank (PDBid: 6f3k).

## Acknowledgements

This work was supported by the European Research Council (StG-2012-311318-ProtDyn2Function to P. S. and CoG-2010-260887 to J. B.) and used the platforms (NMR, EM, isotope-labeling) of the Grenoble Instruct-ERIC Center (ISBG; UMS 3518 CNRS-CEA-UJF-EMBL) with support from FRISBI (ANR-10-INSB-05-02) and GRAL (ANR-10-LABX-49-01) within the Grenoble Partnership for Structural Biology (PSB). The electron microscopy facility is supported by the Rhone-Alpes Region, the Fondation Recherche Medicale (FRM), the fonds FEDER and the GIS-Infrastrutures en Biologie Sante et Agronomie (IBISA). Financial support from the TGIR-RMN-THC Fr3050 CNRS for conducting the research at the 1 GHz spectrometer at the CRMN Lyon is gratefully acknowledged. J.-P.C. acknowledges support by the Agence Nationale de la Recherche (ANR-17-CE11-0018-01).

## Supporting Information

**Figure S1:**
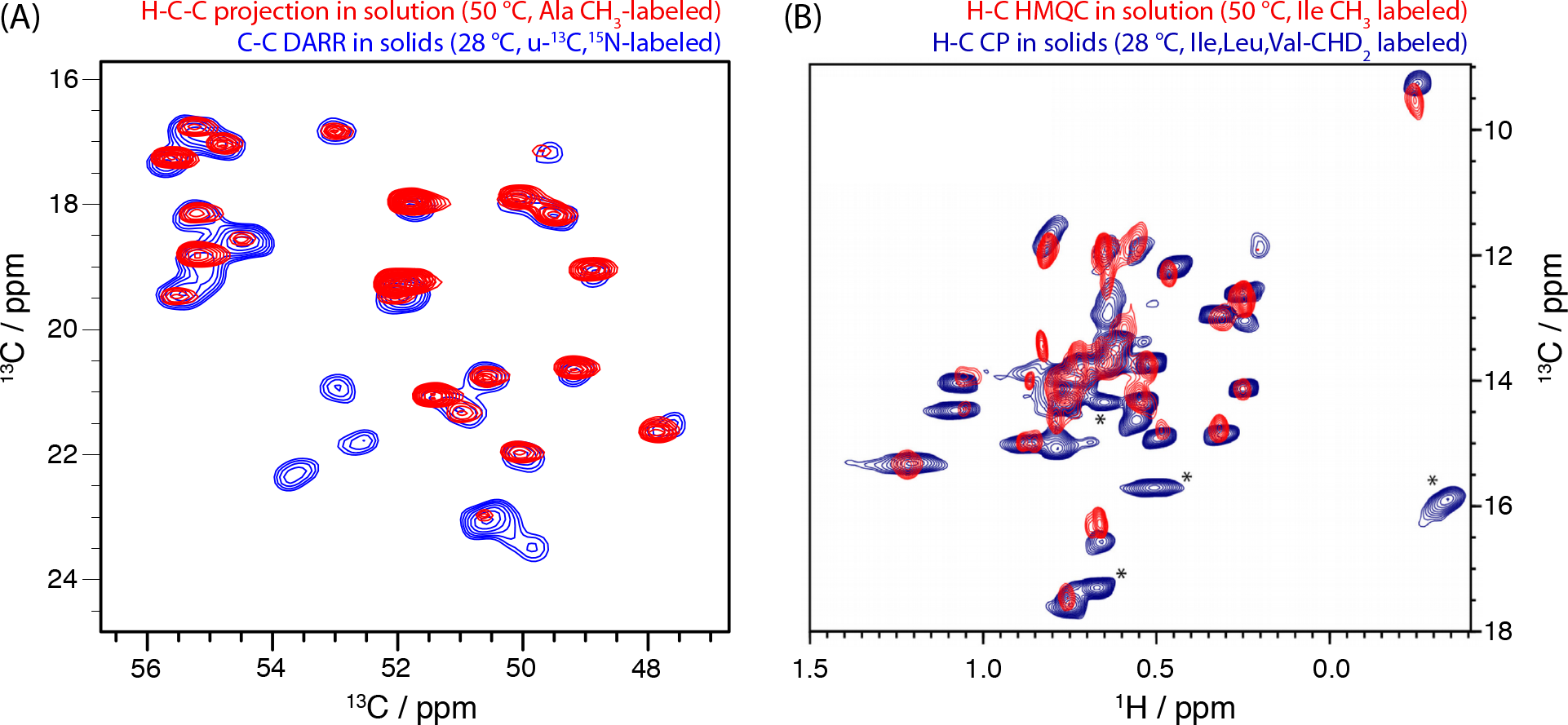
Comparison of solution and MAS NMR spectra of TET2. (A) Alanine C*α*-C*β* correlation spectrum. The solution-state NMR spectrum was collected with a deuterated sample labeled with ^13^C^1^ H_3_ group and a ^13^C spin at the Ca position. The spectrum is a ^1^H detected ^1^H-^13^C-^13^C correlation, projected along the ^1^H dimension. The MAS NMR spectrum is a DARR experiment, recorded on uniformly ^13^C,^15^N labeled sedimented sample, collected at 15 kHz MAS (14.1 T). Note that because of spin relaxation during the transfer steps in the solution-NMR 3D correlation spectrum, several peaks are undetected. The observable peaks match very well the ones in the MAS NMR spectrum. (B) Comparison of ^1^H-^13^C correlation spectra of Ile methyl groups in solution and in an MPD-precipitated TET2 sample. The peaks denoted with an asterisk are resonances from valine sites, which are methyl-labeled in the MAS NMR sample, but not in the solution-NMR sample.

**Figure S2:**
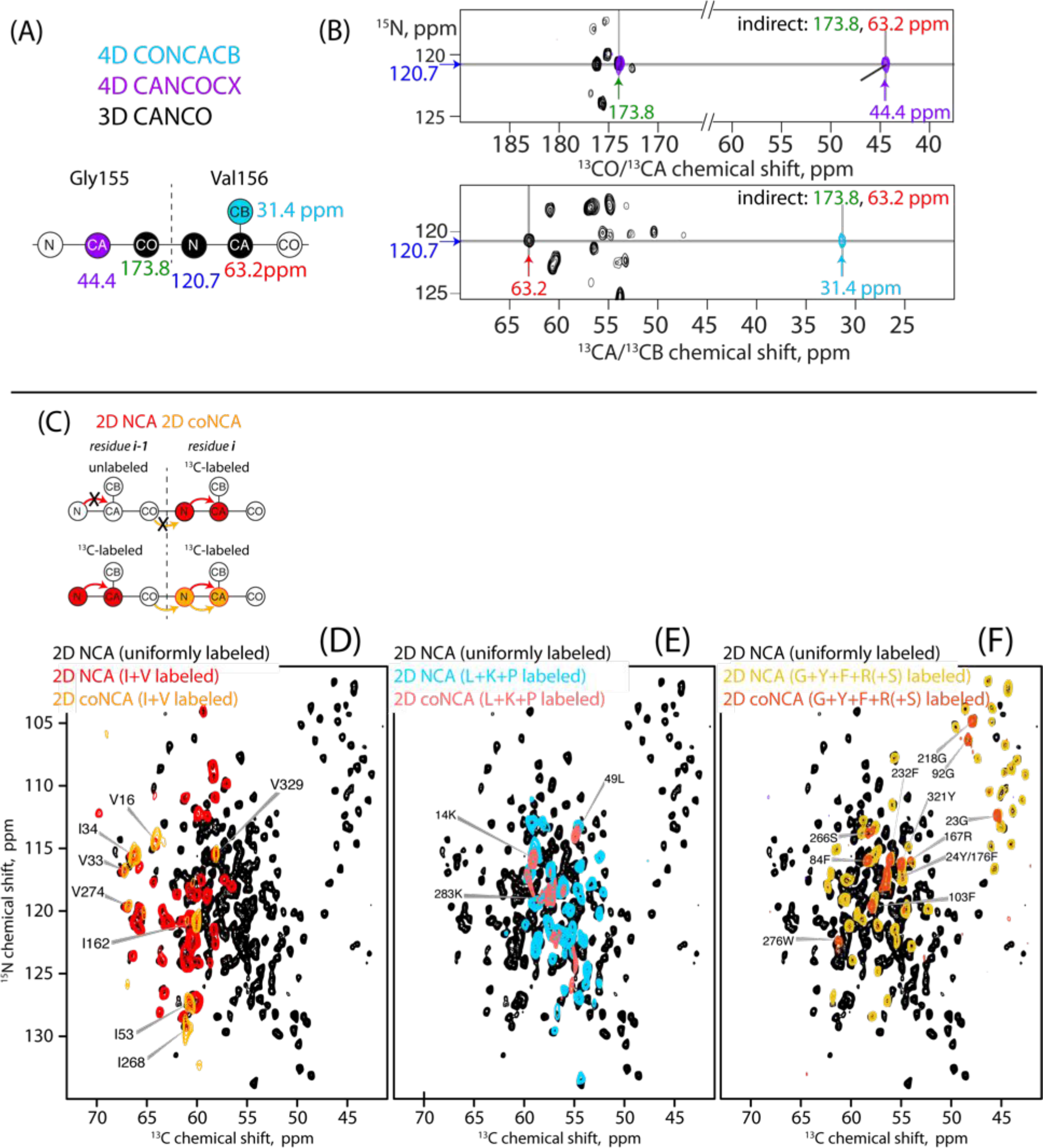
Approach for MAS NMR assignment of TET2 using high-dimensional experiments on uniformly ^13^C,^15^N-labeled samples (A,B) and with the aid of specifically labeled samples (C-F). (A) Schematic coherence transfer and (B) exemplary spectra showing the assignment strategy based on 4D backbone correlation experiments. The CO_*i*−1_, N_*i*_, CA_*i*_ frequencies observed in the CANCO experiment (black) are well resolved and serve as an unambiguous “root peak” for identifying each site. A fourth frequency dimension is added either towards residue *i−1* (CANCOCX) or towards residue *i* (CONCACB), thus allowing to unambiguously correlate five frequencies for each residue pair. (C) Principle of the specific labeling strategy combined with ^15^N-^13^C*α* correlation spectra. In the three different samples only certain types of amino acids are labeled with ^13^C, and the protein is uniformly ^15^N-labeled. In NCA correlation spectra, only residues with ^13^C labeling are observable. In co-N-CA correlation spectra, where the initial magnetization is emanating from the carbonyl of the preceding residue, signals are observed only if two consecutive residues are labeled. In (D)-(F) NCA and coNCA correlation spectra from the three differently labeled samples are presented in colors, and compared to a NCA spectrum of a uniformly ^13^C,^15^N-labeled sample. Note that in the G+Y+F+R sample, residual labeling of serine is also obtained through scrambling from glycine. The spectrum of uniformly ^13^C,^15^N-labeled TET was collected at 1 GHz ^1^H Larmor frequency) while all other spectra were recorded at 600 MHz ^1^H Larmor frequency. Residues highlighted with their sequential number are those observed in the coNCA spectra, i.e., for which the preciding residue is also ^13^C-labeled. A fully annotated NCA spectrum and amide ^1^H,^15^N and methyl ^1^H,^13^C spectra are displayed in Figure S3.

**Figure S3:**
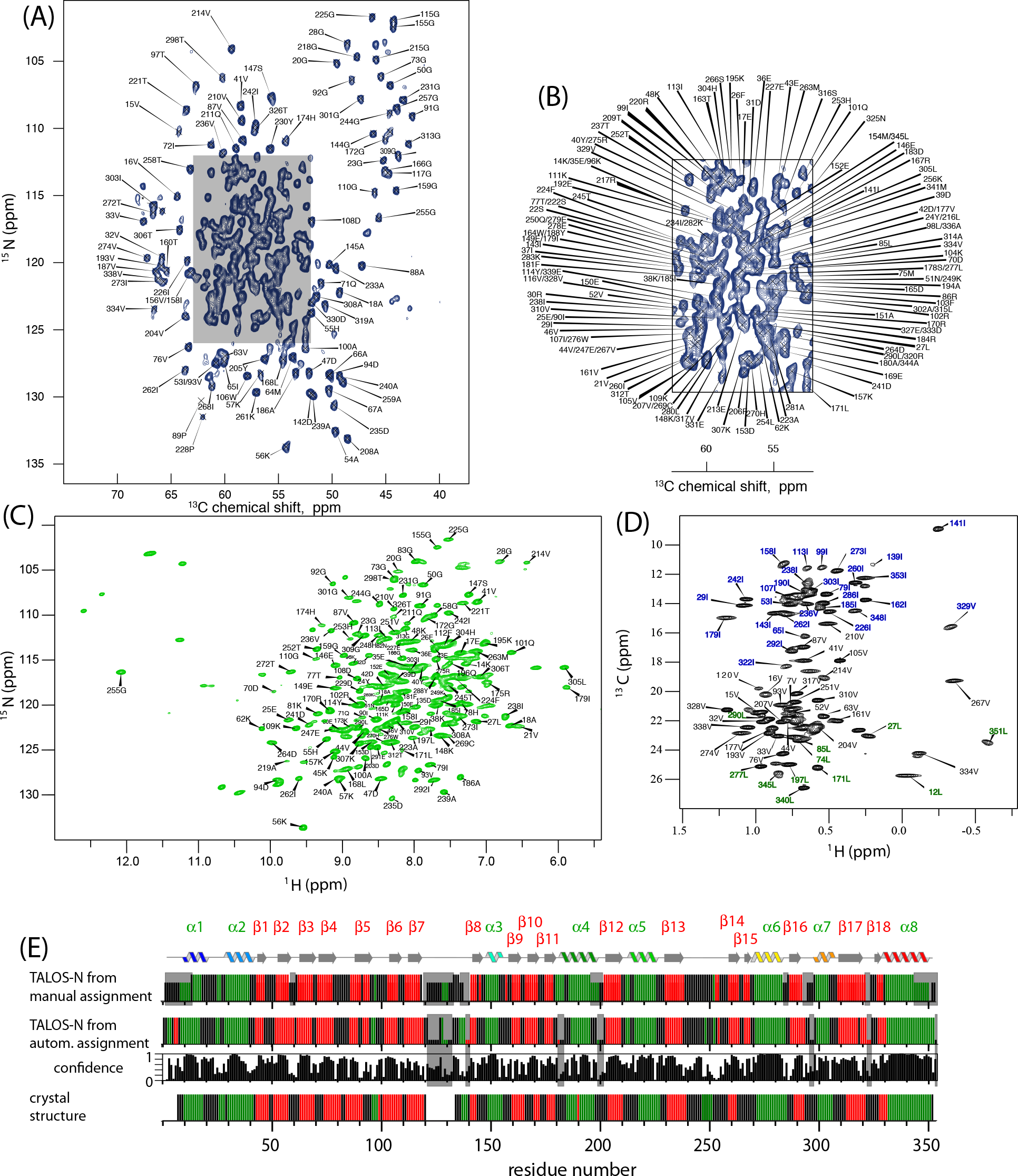
Assigned 2D MAS NMR correlation spectra of TET2. (A) NCA correlation spectrum (^13^C-detected) of uniformly ^13^C,^15^N-labeled TET2 recorded at 1000 MHz ^1^H Larmor frequency, and a zoom of the central part (B). (C) ^1^ H-detected amide ^1^H-^15^N correlation spectrum of uniformly ^2^H,^13^C,^15^N-labeled TET2. (D) ^1^H-detected methyl ^1^H-^13^C MAS NMR correlation spectrum of u-[^2^H,^15^N]-Ile^*δ*1^, Leu^proS^,Val^proS^-[^13^CHD_2_]-labeled TET2 at 600 MHz ^1^H Larmor frequency (14.1 T). (The pro-S position is also referred to as *γ*2 (in Val) and *δ*2 (in Leu).) Ile, Val and Leu methyl groups are annotated in blue, black and green, respectively. (E) Secondary structure in TET2 derived from MAS NMR chemical shift assignments using the program TALOS-N^20^, and comparison to secondary structures in the crystal structure. Loop/coil regions are shown in black, *α*-helices in green and *β*-strands in red. Grey bars in the background denote residues for which no backbone assignment was obtained, comprising the long flexible loop from residues 120-133. Note that for residues where no/insufficient chemical-shift assignments are available, TALOS-N uses a database approach to propose secondary structures; these residues are shown as shorter bars in the TALOS-N plot. For comparison, the results from the manual and automatic (FLYA) assignments are shown. The confidence level of the TALOS-N secondary structure predicting is shown for the FLYA data set. The lowest shows the secondary structures as obtained from the crystal structure (PDB entry 1Y0R) and the DSS algorithm, determined with pymol. Missing bars denote residues not modeled in the crystal structure.

**Figure S4:**
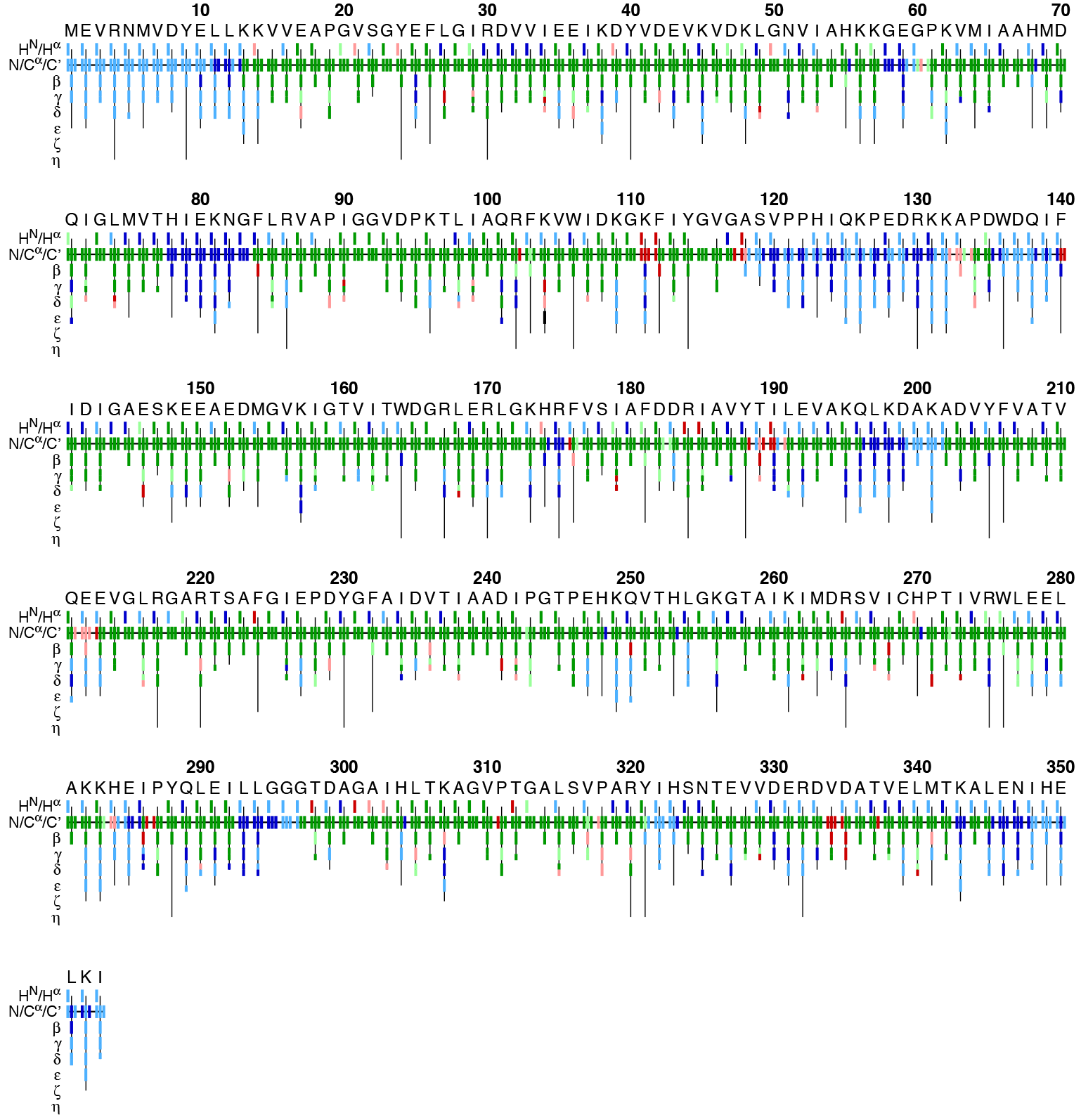
Automatic assignment of MAS NMR spectra of TET2 using the program FLYA.^19^ The program used the following spectra for assignment: NCA, NCO, NCOCX, CANCO, NCACX, NCACB, NcoCACB, CONCACB, CANCOCX, NCOCX(GRYF), NCOCX(ILV), NCOCX(LKP), NCACX(GRYF), NCACX(ILV), NCACX(LKP), hCANH, hCONH, hcoCAcoNH, CCCdarrdarr(ILV), CCC, caNCO(GRYF), coNCA(GRYF), coNCA(ILV), coNCA(LKP). The color code reflects the assignment for each atom, and compares it to the manual assignment, as follows. Green: Automatic and manual assignment agree; red: automatic assignment differs from manual assignment; blue: no manual assignment available. Dark colors (green, blue, red) denote atoms for which the FLYA assignment has converged, i.e., the same assignment is found in more than 80% of the 20 independent runs, while light colors denote atoms for which FLYA has not converged, i.e., the results have less confidence. The row labeled H^N^/H^*α*^ shows for each residue HN on the left. The N/C/C*α* row shows for each residue the N, C*α*, and C’ assignments from left to right. The rows labeled with *β*-*η* show the side-chain assignments for the heavy atoms. In the case of branched side chains, e.g. Val and Leu, the corresponding row is split into an upper part for one branch and a lower part for the other branch.

**Figure S5:**
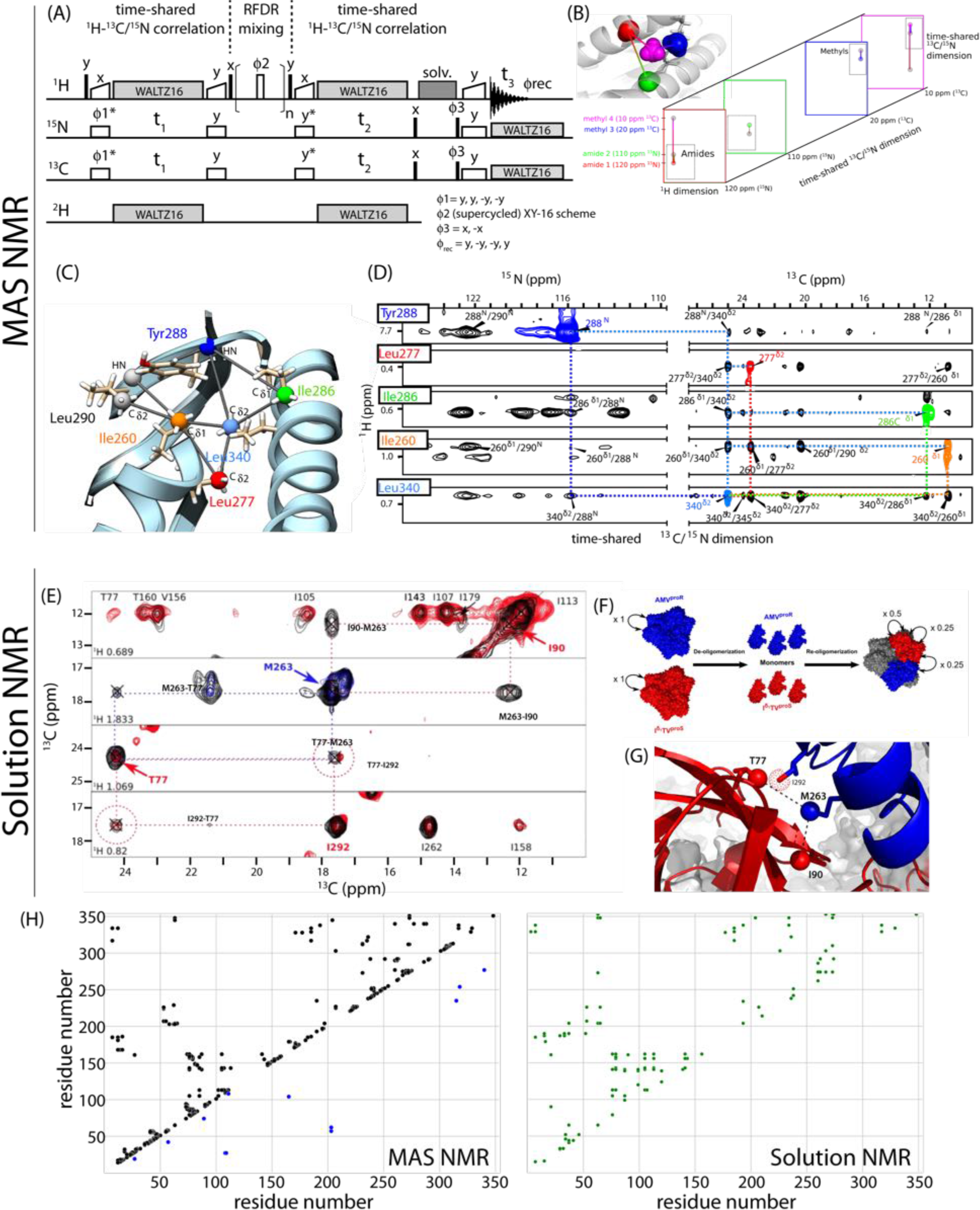
Distance restraints from MAS NMR and solution-NMR experiments. (A) Pulse sequence used for the simultaneous measurement of ^1^H-^1^H spatial proximities between amide and methyl moieties under fast-MAS conditions, for application to deuterated, amide-protonated and methyl ^13^CHD_2_ labeled samples. Filled and open narrow rectangles denote 90° and 180° pulses, respectively, and wide open rectangles denote cross-polarization steps, applied with a linear ramp on the ^1^H channel. The RFDR mixing, of a total duration of 5 ms, is comprised of a train of *n* equally-spaced pulses where the centers of two consecutive pulses are separated by the MAS rotor period; the pulse phase is alternated according to the XY16 scheme. The cross-polarization steps simultaneously transfer polarization from amide-^1^H to ^15^N and from the methyl-^1^H to the methyl-^13^C (out and back). Pulse phases are denoted above the respective symbols. Quadrature detection in the indirect dimensions (t_1_, t_2_) was performed by varying the phase of the ^13^C and ^15^N cross-polarization irradiation prior to the chemical-shift (denoted by an asterisk) according to the States-TPPI scheme. Decoupling of spin-spin couplings during direct and indirection dimensions and the solvent-suppression scheme (”solv.”) are denoted by grey rectangles. (B) Schematic representation of the outcome of this experiment, showing connections between two amide sites, between two methyl sites and between an amide and a methyl site. (C) Experimental distance restraints from the RFDR experiment shown on part of the TET2 structure. (D) Excerpts from the 3D RFDR MAS NMR spectrum, showing the cross-peaks supporting the distance restraints shown in (C). (E) Subunit-specific labeling of TET2 to filter inter-subunit distance information. Example strips from 3D HMQC-NOESY-HMQC spectra recorded on either u-[^2^H,^15^N],Ile^*δ*1^, Thr^*γ*2^,Val^proS^-[^13^CH_3_] (red), u-[^2^H,^15^N],Met^*ϵ*^,Ala^*β*^,Val^proR^-[^13^CH_3_] (blue) or a sample in which these two differently labeled subunit types were mixed in a 1:1 ratio (black). (F) Labeling scheme for these spectra. The numbers reflect the relative cross-peak intensities resulting from the stochastic mixing scheme. (G) Illustration of inter-subunit distance restraints detected between M263 in one subunit and T77 and I90 in another subunit. An additional methyl group from which an intra-subunit contact is observed in the spectra in (C) is shown with a dotted sphere. Note that the inter-subunit distance restraints have not been used directly in this work, but we have only used this information for excluding inter-subunit restraints from the calculations (see Methods). A total of only eight inter-subunit distances has been identified, and this set of experiments might not be crucial in the presented approach. (H) Residue contact maps showing the restraints from the ^1^H-^1^H MAS NMR RFDR experiment connecting amide and methyl sites (black), the ^13^C DARR experiment on LKP-labeled TET2 (blue), and the solution-state NOESY experiments (green). The eight inter-subunit restraints have been removed from this graph.

**Figure S6:**
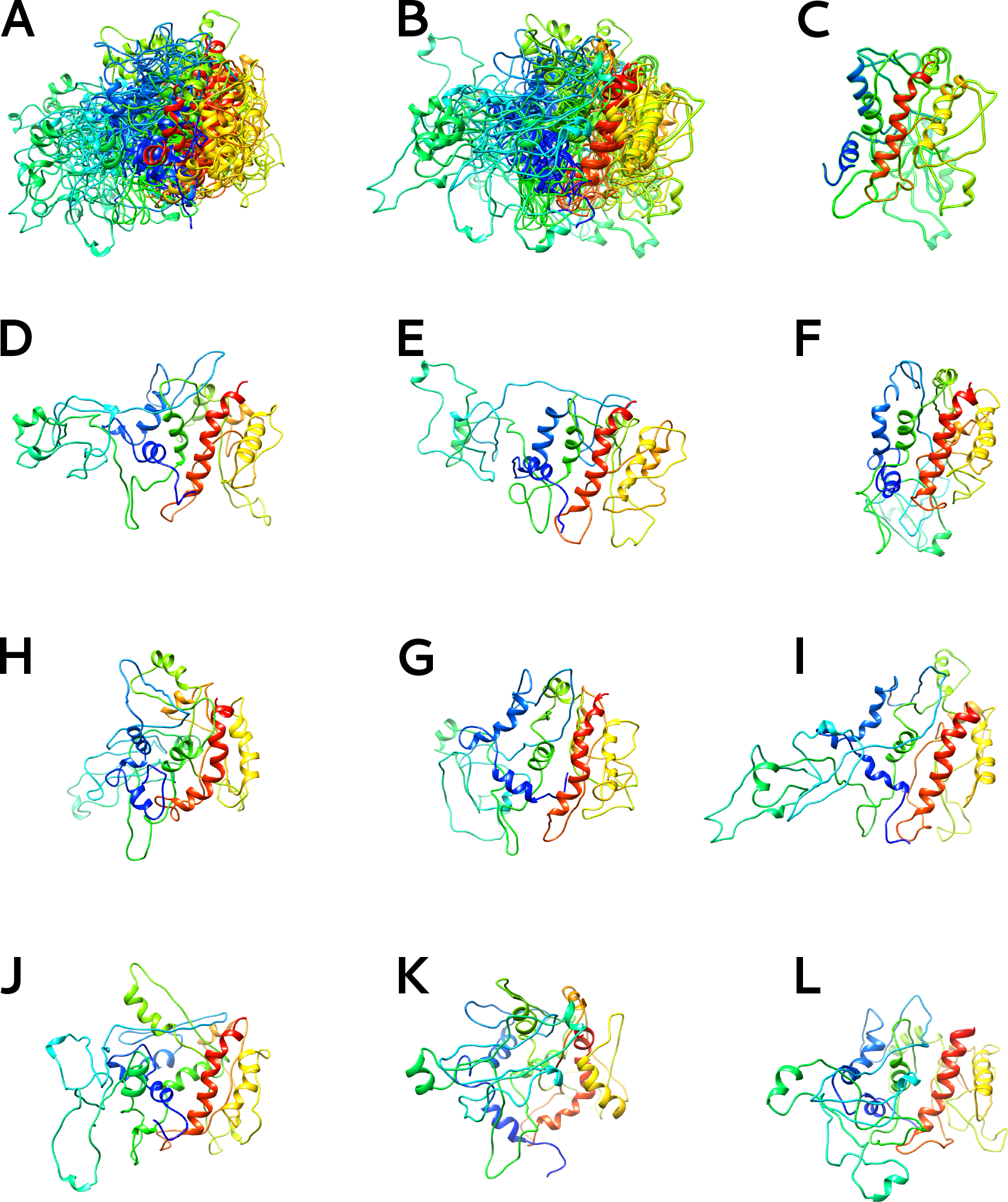
Structure calculation of the monomer of TET2 using only the liquid and solid-state NMR derived restraints. Figure (A) represents the NMR ensemble of 10 structures superimposed on their secondary structure elements (backbone RMSD: 10.5 ± 1.6 Å). Figure (B) shows the same NMR structure ensemble superimposed on the largest helix *α*8, each structure of this ensemble are displayed individually on figures (C) to (L).

**Figure S7:**
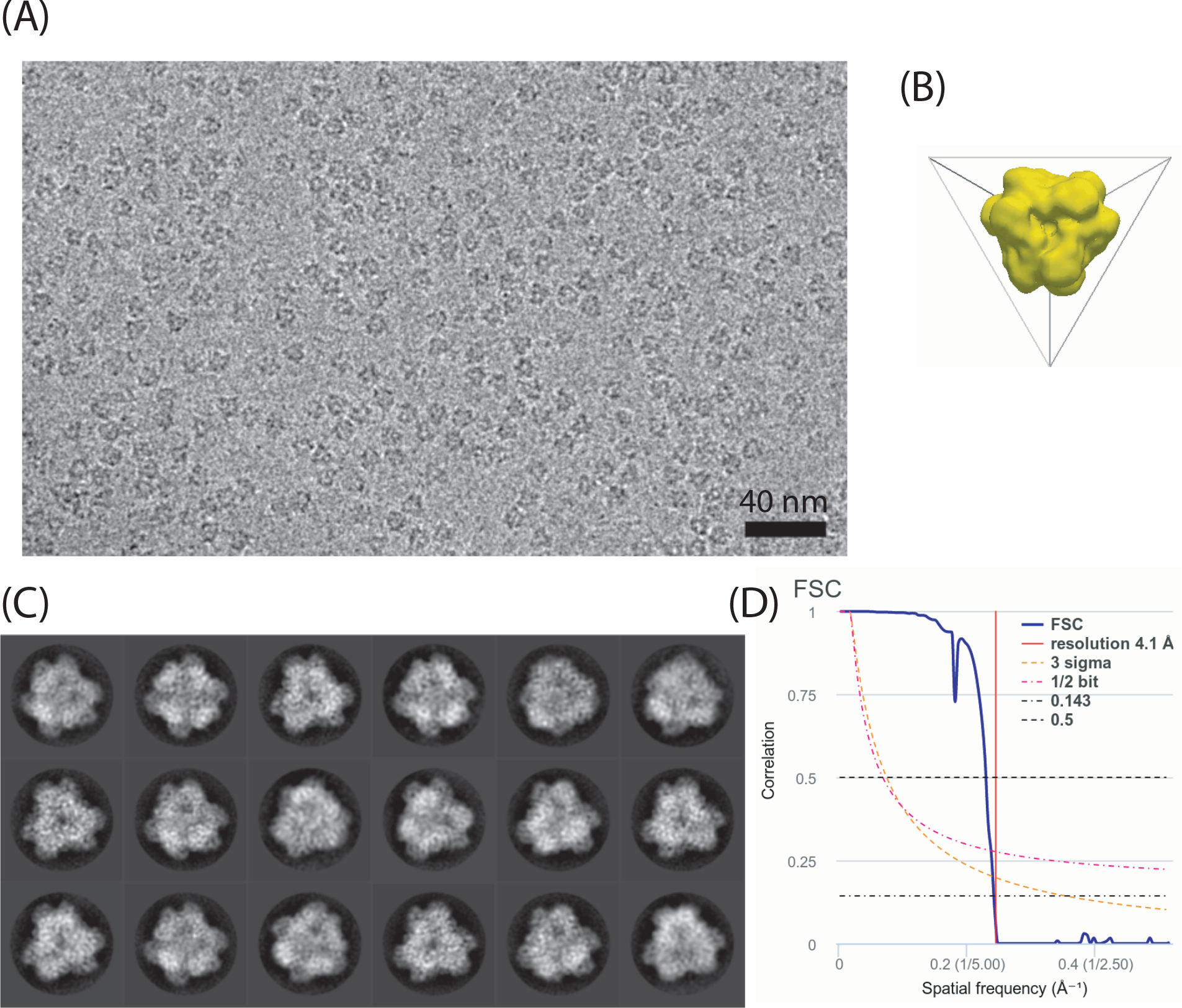
(A) Typical cryo-electron micrograph showing TET particles (darker than the background densities) in random orientations. (B) Used initial low resolution 3D model obtained by imposition of tetrahedral symmetry. (C) Best eighteen 2D class averages obtained from 27,130 particles. (D) Fourier Shell Correlation plot for the presented 3D cryo-EM reconstruction.

**Figure S8:**
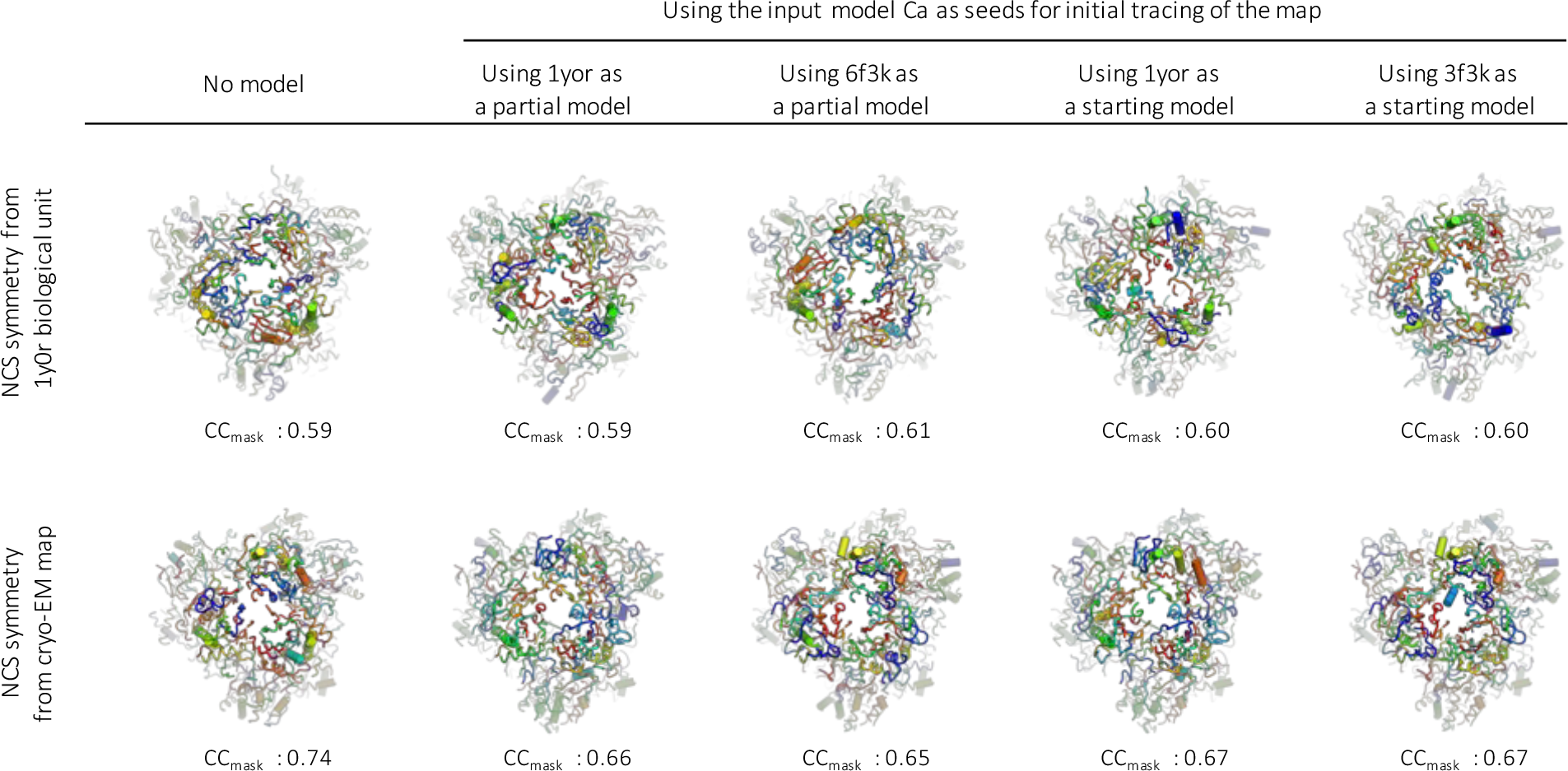
Attempts to determine the structure from the EM map only. Direct rebuilding from the 4.1 Å cryo-EM map fails to produce interpretable structures, regardless of the symmetry that is imposed (or not) for model-building, and of whether (or not) a partial or a starting model is provided. Briefly, we used phenix.map_to_model (10 rebuild cycles) to attempt rebuilding the structure directly from the cryo-EM map (4.1 Å resolution) using either no prior information (no model), or the X-ray (lyOr) or the NMR+cryo-EM (6f3k, this paper) structures as starting coordinates. We tested inclusion of these models either as “partial” or “starting” models, and in all cases, imposed the use of the input-model C*α*-atoms as seeds for the initial tracing of the map. Rebuilding was attempted with (upper row) or without (lower row) imposing the known symmetiy of the particle. In both case, phenix.map_to_model was able to trace the map, but detection of the NCS symmetry from the map (lower panel), as opposed to inferred from the X-ray pdb structure (upper panel), drastically improved the quality of the model. At the map resolution, however, phenix.modeLtojnap was not able to interpret the electron density in terms of secondary structure and/or sequence at least not in 10 rebuild cycles. Arguably, the traced model produced by phenix.map_to_model would have been interpretable by an experienced crystallographer or electron microscopist, but reaching the quality of 6f3k would have been veiy challenging.

**Figure S9:**
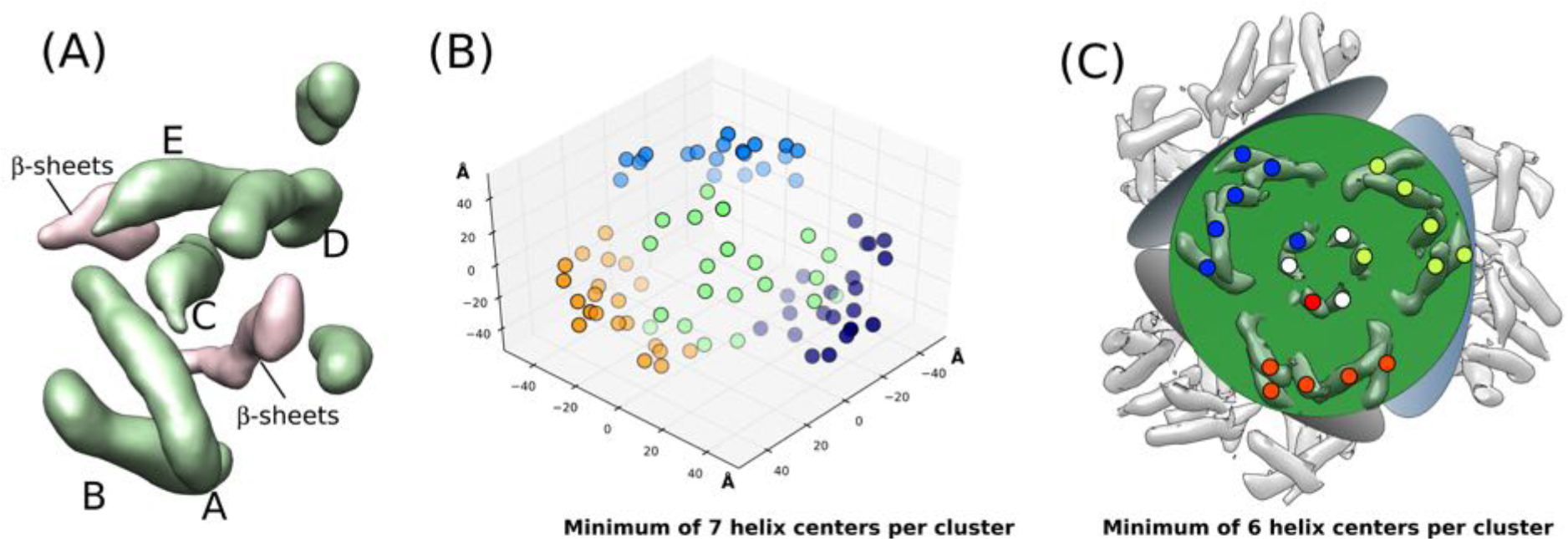
Automatic detection and clustering of secondary structures. (A) Secondary structures detected by *gorgon* ^47^ from the 8 Å EM map of TET2, zooming on one TET2 subunit. *α*-helices and *β*-sheet regions are depicted in green and pink, respectively. Note that in this study we have only used five *α*-helices, labeled with A to E, and we have not used the information about the location of the *β*-sheets. Including these regions is straightforward and may prove useful in other applications, with proteins containing primarily *β*-sheets. (B,C) Cluster analysis of the *α*-helices from the EM map, aimed at identifying the *α*-helices belonging to the same monomer. This automatic distance-based clustering analysis used the python library sklearn.cluster, and needs as input (i) the coordinates of the points in space to be clustered – here, the centers of the 84 automatically detected *α*-helices –, (ii) the minimum number of elements in each cluster and (iii) the maximum distance between the points belonging to a given cluster. Note that no information about the symmetry or the number of subunits (number of clusters) is provided. In (B), the minimum number of elements belonging to the same cluster was set to 7. The points distributed in space, representing the centers of the *α*-helices in the dodecamer, are color-coded according to the group/cluster to which they are assigned. While this clustering does not retrieve the monomer subunits, it detects clusters of 21 points, which correspond to trimers of TET2. In (C), this analysis was repeated, setting the minimum number of elements per cluster to 6. In this case, the six points assigned to each cluster (shown in blue, red or green) indeed correspond to helix centers of a given subunit. The additional white dots represent the helix centers that could not be assigned to any cluster using the distance-based clustering algorithm. The green plane is shown for facilitating visualization of one trimer within the dodecamer. In Step 1 of our structure calculation protocol we have used the five *α*-helices within these clusters which are closest in space to each other, omitting the sixth helix.

**Figure S10:**
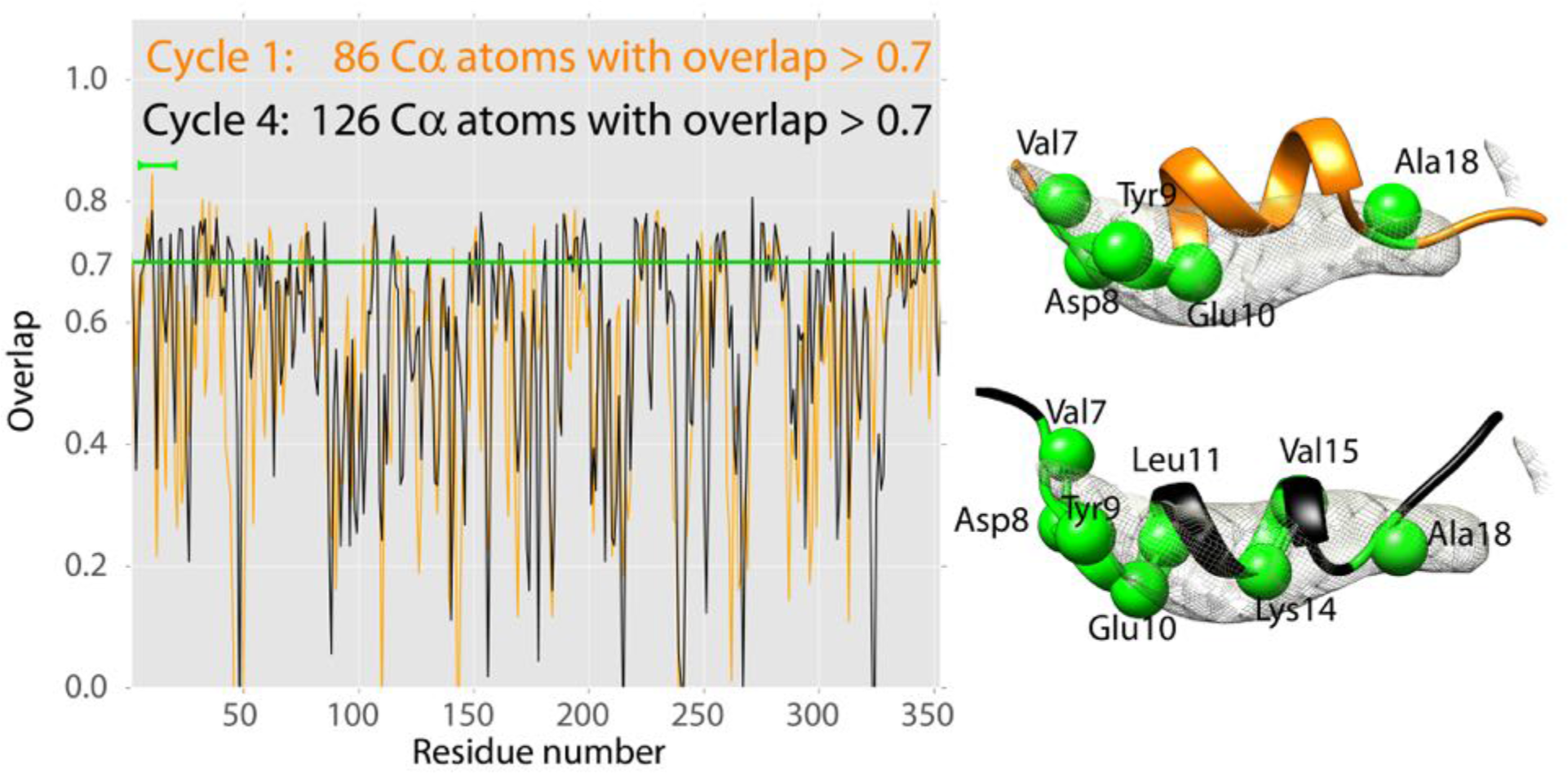
Illustration of the procedure through which, in Step 2 of this approach, residues were restrained to real-space coordinates, based on their overlap with the electron density map. Briefly, after flexible fitting with iModFit the structure was compared to the 8 Å electron density map to evaluate how well different parts of the chain reside inside the density. To this end, the *molmap* and *measure correlation* modules in UCSF Chimera were used to compute the overlap of an in silico map, constructued from the backbone atoms of a given residue, with the experimental electron density. The correlation between these two maps, a value ranging from 1 (atom is fully inside density) to 0 (atom outside density) was used as a criterion whether the residue resides within the map or not. If the value was above 0.7 for a given residue, the Cα atom of this residue was then kept in place in the subsequent CYANA calculation by adding restraints from this atom to the other atoms that are fixed in space (using a tolerance of 0.5 Å.) (A) Residue-wise plot of the overlap score in the first and fourth iteration of Step 2. The green bar on the top left indicates the residues which are shown in (B). (B) Example of the overlap in the first and fourth iteration. C*α* atoms with an overlap score above 0.7 are indicated with green spheres and were constrained in the space with a tolerance of ±2*Å*.

**Figure S11:**
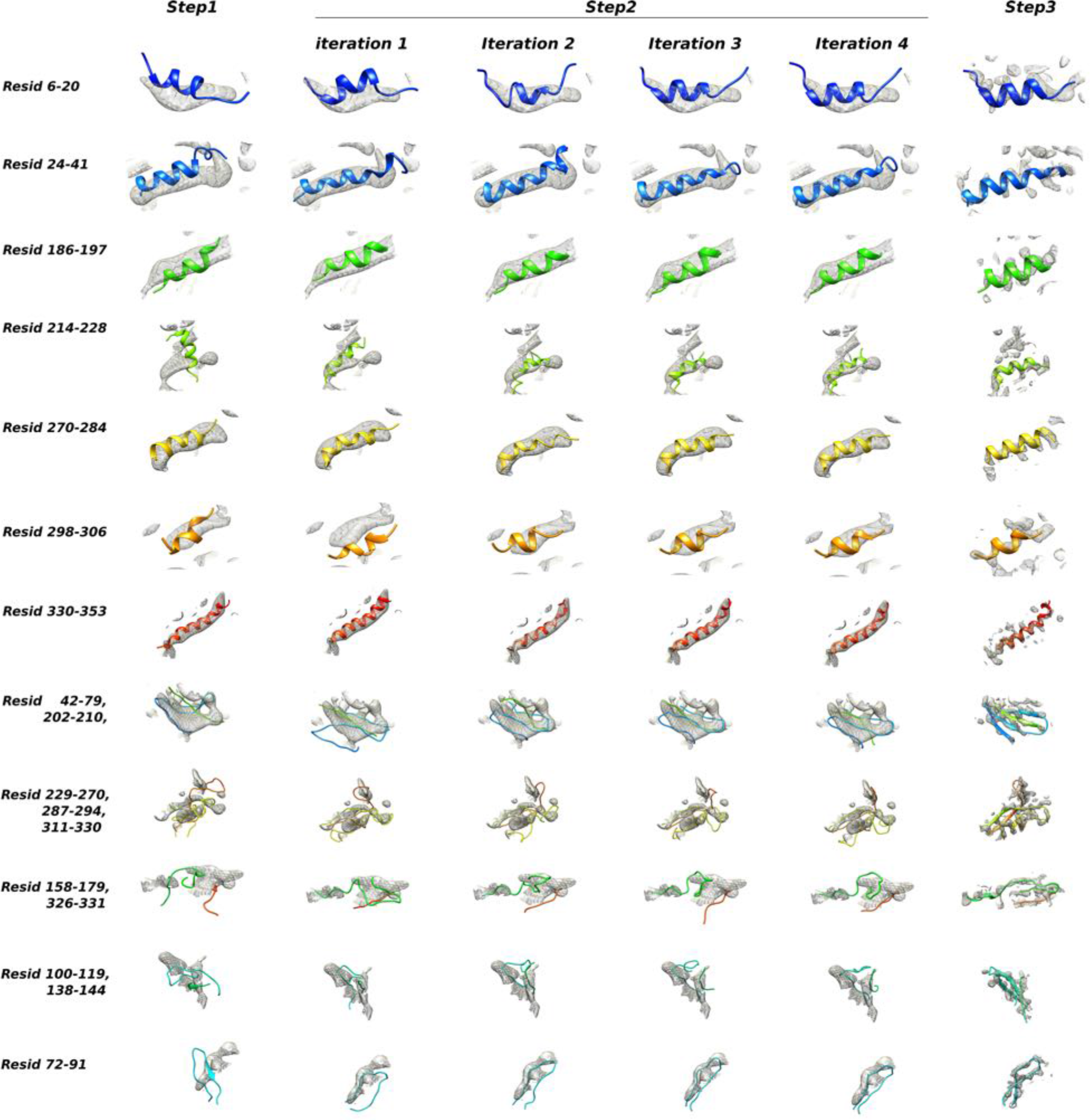
Evolution of the structure of TET2 along the various steps of the integrated NMR/EM structure calculation approach, denoted as Step 1 to 3 in Figure 2. The structures shown correspond to the lowest-energy models generated by CYANA (Steps 1 and 2) or XPLOR-NIH (Step 3).

**Table S1.** *(Preceding page)* Details about the helix-to-density assignments. The number of helix-to-density assignments for eight *α*-helices detected in the sequence, placed into the seven *α*-helical densities detected in the EM map (i.e., leaving out one helix in each combination), ignoring the helix polarity, can be calculated as 8! = 40320. Including the helix polarity increases this number to 8! × 2^7^ ≈ 5 · 10^7^. Considering the possibility that the seven densities may not be ascribed unambiguously to a given subunit further increases this number. We solved this latter point (identifying densities of one subunit) by a cluster analysis (see Figure S9) and retaining only the five longest helical densities, which are also closest in space. We explicitly disregarded the helix polarity, by using only the center of the helices in the CYANA calculation, letting the rotation free. The number is furthermore greatly reduced by comparing the lengths of *α*-helices along the sequence with the helical densities. (A) This table shows the lengths (middle column) of the helical densities from the EM map (denoted as A-E as shown in Figure 1G, left column), and the *α*-helices along the sequence (right column, see Figure 1D) that may match the lengths of these densities. Hereby, we have considered that a helix has a length of 1.4 Å per residue. In selecting the possible helix-to-density assignments, we applied a tolerance of ±2 amino acids (corresponding to ±2.8 Å). As a result, only 1 × 2 × 5 × 4 = 40 helix-to-density assignments remain that are checked by structure calculations. (B) Identity of the residues assumed to be in the centers of the *α*-helices, as obtained from the NMR assignments and TALOS-N. The lengths of the helices in Å was estimated from the length in number of residues, as obtained by TALOS-N, assuming a canonical translation length per residue of ca. 1.45 Å. (C) Practical implementation of the helix-helix distance restraints, shown here for the correct helix-to-density assignment. These distance restraints were used in CYANA, alongside the other NMR distance restraints and backbone dihedral-angle restraints.

| (A) $\alpha$ helices cryo-EM | Length (cryo-EM) | $\alpha$ helices NMR |
| --- | --- | --- |
| A | 30.8 Å | $\alpha 8$ |
| B | 20.2 Å | $\alpha 6, \alpha 4$ |
| C | 21.7 Å | $\alpha 6, \alpha 4$ |
| D | 18.1 Å | $\alpha 6, \alpha 4, \alpha 5, \alpha 2, \alpha 1, \alpha 7, \alpha 3$ |
| E | 13.1 Å | $\alpha 5, \alpha 2, \alpha 1, \alpha 7, \alpha 3$ |
| (B) Residues placed in centers of the α-helices and estimated lengths in Å | | |
| $\alpha 1$ | Lys 13 | 13.0 Å |
| $\alpha 2$ | Ile 34 | 13.0 Å |
| $\alpha 3$ | Ala 151 | 8.6 Å |
| $\alpha 4$ | Leu 191 | 15.8 Å |
| $\alpha 5$ | Arg 220 | 13.0 Å |
| $\alpha 6$ | Leu 277 | 18.7 Å |
| $\alpha 7$ | Ile 303 | 8.6 Å |
| $\alpha 8$ | Met 341 | 31.7 Å |
| (C) Distance restraints corresponding to the correct helix-to-density assignment |  |  |
| Met 341 Cα | Leu 277 Cα | $10.84 \pm 0.50$ Å |
| Met 341 Cα | Leu 191 Cα | $10.46 \pm 0.50$ Å |
| Met 341 Cα | Ile 34 Cα | $20.36 \pm 0.50$ Å |
| Met 341 Cα | Lys 13 Cα | $19.24 \pm 0.50$ Å |
| Leu 277 Cα | Leu 191 Cα | $18.30 \pm 0.50$ Å |
| Leu 277 Cα | Ile 34 Cα | $28.11 \pm 0.50$ Å |
| Leu 277 Cα | Lys 13 Cα | $28.92 \pm 0.50$ Å |
| Leu 191 Cα | Ile 34 Cα | $10.43 \pm 0.50$ Å |
| Leu 191 Cα | Lys 13 Cα | $12.49 \pm 0.50$ Å |
| Ile 34 Cα | Lys 13 Cα | $12.52 \pm 0.50$ Å |

**Table S2.**
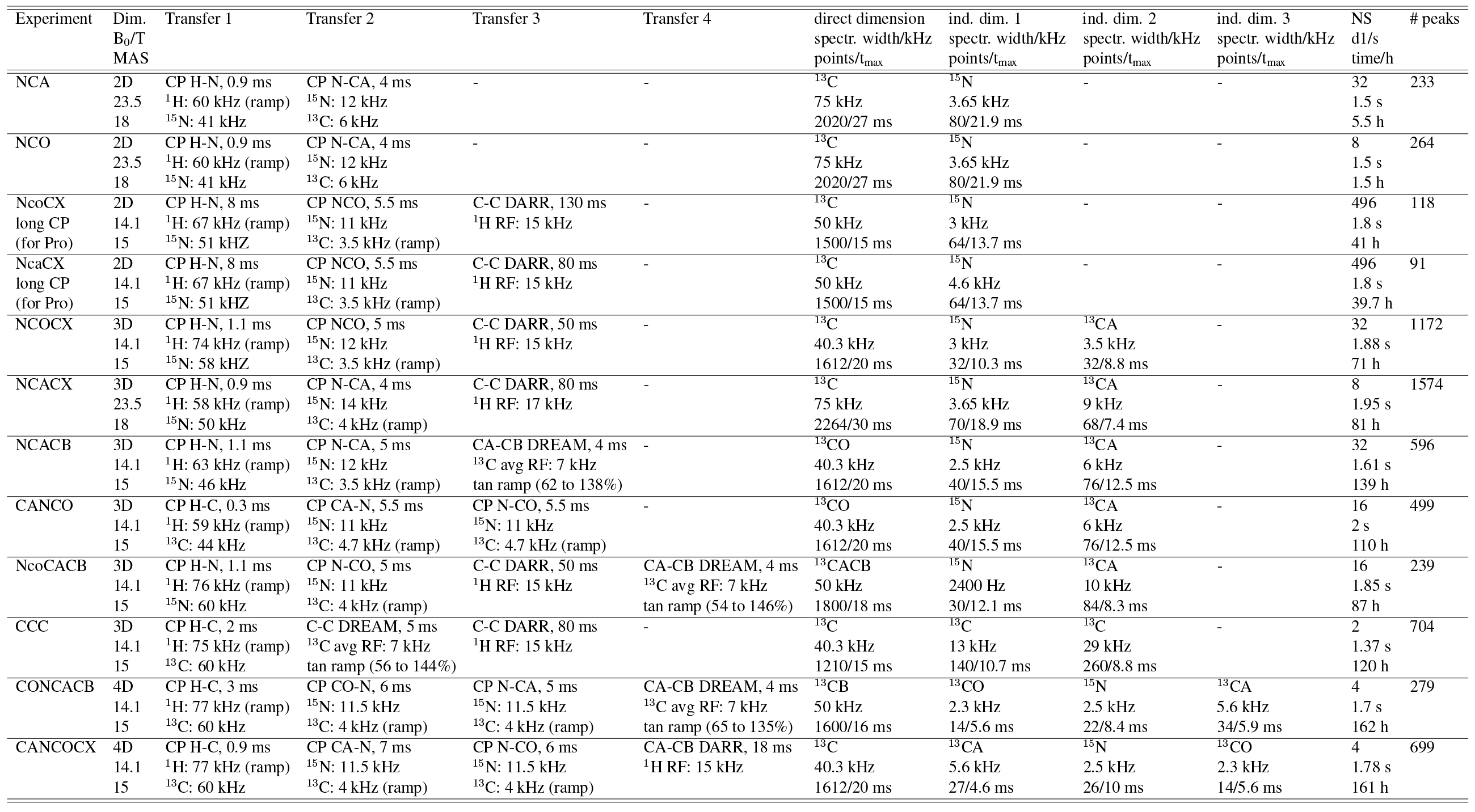
Acquisition parameters of MAS NMR spectra for resonance assignment of protonated, ^13^C,^15^N labeled TET2 in 3.2 mm rotors. CP transfer steps between ^15^N and ^13^CA or ^13^CO used a ^13^C RF carrier either at the C^*α*^ (54 ppm) or CO (175 ppm) frequency. The MAS frequency is denoted in kHz. All spectra at 14.1 T were acquired on a Agilent spectrometer; the three spectra (two 2Ds and one 3D) at 23.5 T were recorded on a Bruker Avance spectrometer. The two experiments denoted with “long CP” used a longer ^1^H-^15^N cross-polarization to promote proline signals. The column “peaks” is the number of detected peaks used for the automatic assignment in FLYA.

**Table S3.**
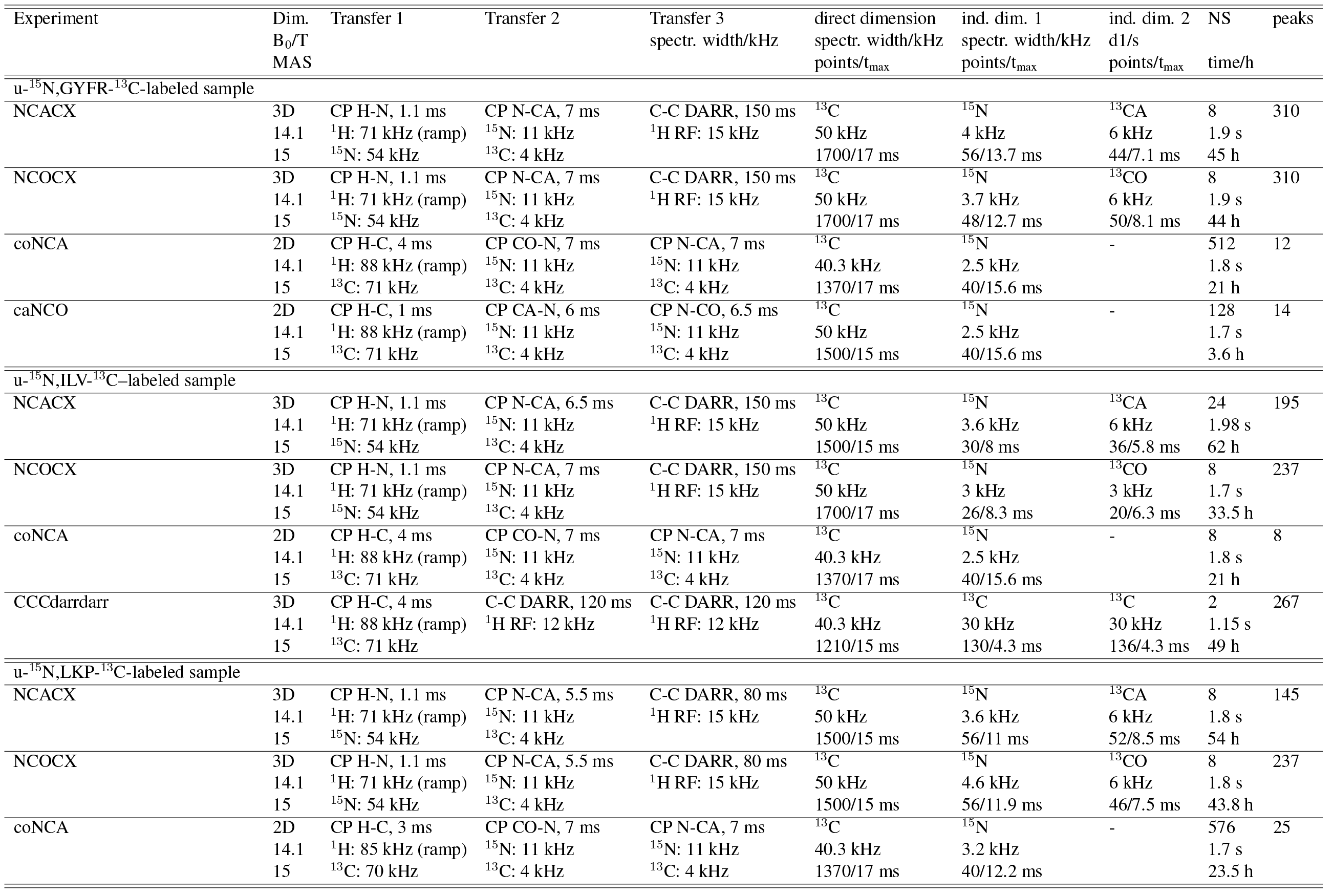
Acquisition parameters of MAS NMR spectra for resonance assignment from specifically labeled samples. Spectra were recorded on a 600 MHz Agilent spectrometer in 3.2 mm rotors. Three samples were used, labeled uniformly with ^15^N by addition of ^15^NH_4_ to the growth medium, and labeled specifically with the specified amino acids (specified in the table), by addition of these amino acids with u-^15^N,^13^C labeling. An exception is the labeling of Leu and Val: here, not the amino acids but the ^13^C-labeled acetolactate was added^40^; as a consequence, ^13^C is incorporated at the C^α^ position of Val, but not of Leu, and Leu residues are unobservable in all experiments going through C^*α*^.

**Table S4.** Acquisition parameters of MAS NMR spectra for resonance assignment using proton-detected experiments on u-[^2^H,^13^C,^15^N]-labeled sample, back-exchanged to 100% in H_2_0-buffer. The hcaCBcaNH experiment has not been used for the automatic assignment, but only collected afterwards to confirm assignments manually

| Experiment | Dim.<br>$B_0/T$<br>MAS | Transfer 1 | Transfer 2 | Transfer 3 | Transfer 4 | direct dimension<br>spectr. width/kHz<br>points/$t_{\text{max}}$ | ind. dim. 1<br>spectr. width/kHz<br>points/$t_{\text{max}}$ | ind. dim. 2<br>spectr. width/kHz<br>points/$t_{\text{max}}$ | NS<br>d1/s<br>time/h | peaks |
| --- | --- | --- | --- | --- | --- | --- | --- | --- | --- | --- |
| hNH | 2D | CP H-N, 1 ms | CP N-H, 1 ms | - | - | $^1\text{H}$ | $^{15}\text{N}$ | | 128 | 146 |
| | 14.1<br>55 | $^1\text{H}$: 95 kHz (ramp)<br>$^{15}\text{N}$: 40 kHz | $^1\text{H}$: 95 kHz (ramp)<br>$^{15}\text{N}$: 40 kHz | | | 20.1 kHz<br>1210/30 ms | 3.04 kHz<br>112/36.5 ms | | 0.95 s<br>8 h | |
| hCONH | 3D | CP H-CO, 3 ms | CP CO-N, 6 ms | CP N-H, 1 ms | - | $^1\text{H}$ | $^{15}\text{N}$ | $^{13}\text{CO}$ | 128 | 304 |
| | 14.1<br>38 | $^1\text{H}$: 95 kHz (ramp)<br>$^{15}\text{N}$: 40 kHz | $^{15}\text{N}$: 22 kHz (ramp)<br>$^{13}\text{C}$: 13 kHz | $^1\text{H}$: 95 kHz (ramp)<br>$^{15}\text{N}$: 40 kHz | - | 20.1 kHz<br>1210/30 ms | 2.8 kHz<br>46/16 ms | 3 kHz<br>50/16.3 ms | 0.95 s<br>44 h | |
| hCANH | 3D | CP H-CO, 3 ms | CP CO-N, 6 ms | CP N-H, 1 ms | - | $^1\text{H}$ | $^{15}\text{N}$ | $^{13}\text{CO}$ | 16 | 255 |
| | 14.1<br>38 | $^1\text{H}$: 95 kHz (ramp)<br>$^{15}\text{N}$: 40 kHz | $^{15}\text{N}$: 22 kHz (ramp)<br>$^{13}\text{C}$: 13 kHz | $^1\text{H}$: 95 kHz (ramp)<br>$^{15}\text{N}$: 40 kHz | - | 20.1 kHz<br>1210/30 ms | 2.8 kHz<br>46/16 ms | 3 kHz<br>68/9.6 ms | 1.04 s<br>64 h | |
| hcoCAcoNH | 3D | CP H-CO, 3 ms | CO-CA out/back | CO-N CP | CP N-H, 1 ms | $^1\text{H}$ | $^{15}\text{N}$ | $^{13}\text{CA}$ | 16 | 169 |
| | 14.1<br>38 | $^1\text{H}$: 95 kHz (ramp)<br>$^{15}\text{N}$: 40 kHz | INEPT<br>delay 5 ms | $^{15}\text{N}$: 22 kHz (ramp)<br>$^{13}\text{C}$: 13 kHz | $^1\text{H}$: 95 kHz (ramp)<br>$^{15}\text{N}$: 40 kHz | 20.1 kHz<br>1210/30 ms | 2.3 kHz<br>40/17 ms | 15 kHz<br>106/7.0 ms | 1.1 s<br>92 h | |
| hcaCBcaNH | 3D | CP H-CA, 4 ms | CA-CB out/back | CA-N CP | CP N-H, 1 ms | $^1\text{H}$ | $^{15}\text{N}$ | $^{13}\text{CA}$ | 16 | not |
| | 14.1<br>53 | $^1\text{H}$: 85 kHz (ramp)<br>$^{15}\text{N}$: 33 kHz | INEPT<br>delay 6.5 ms | $^{15}\text{N}$: 38 kHz (ramp)<br>$^{13}\text{C}$: 14 kHz | $^1\text{H}$: 82.5 kHz (ramp)<br>$^{15}\text{N}$: 38 kHz | 20.1 kHz<br>1210/30 ms | 2.55 kHz<br>64/25 ms | 11.3 kHz<br>192/8.4 ms | 1.1 s<br>58 h | used in<br>FLYA |

**Table S5.** Acquisition parameters of experiments used for distance restraint measurements.

| Experiment | Dim.<br>B<sub>0</sub>/T<br>sample | Through-bond<br>transfer | Through-space<br>transfer | direct dimension<br>spectr. width/kHz<br>points/t<sub>max</sub> | ind. dim. 1<br>spectr. width/kHz<br>points/t<sub>max</sub> | ind. dim. 2<br>spectr. width/kHz<br>points/t<sub>max</sub> | NS<br>dl/s<br>time/h |
| --- | --- | --- | --- | --- | --- | --- | --- |
| h(N/C)h-RFDR-h(N/C)-H | 3D | CP H-N & H-C, 2.5 ms | <sup>1</sup>H RFDR, 8 ms (440 cycles) | <sup>1</sup>H | <sup>15</sup>N & <sup>13</sup>C | <sup>15</sup>N & <sup>13</sup>C | 4 |
| MAS NMR | 14.1 | <sup>1</sup>H: 95 kHz (ramp) | 100 kHz <sup>1</sup>H pulses | 12.02 kHz | 6.8 kHz | 6.8 kHz | 0.83 s |
| MAS: 55 kHz | sample 6 | <sup>15</sup>N/<sup>13</sup>C: 41/43 kHz |  | 1200/50 ms | 164/24.1 ms | 168/24.7 ms | 118 h |
| LKP <sup>13</sup>C DARR | 2D | CP H-C, 1.6 ms | <sup>13</sup>C-<sup>13</sup>C DARR, 350 ms | <sup>13</sup>C | <sup>13</sup>C | - | 192 |
| MAS NMR | 14.1 | <sup>1</sup>H: 70 kHz (ramp) | <sup>1</sup>H RF: 12 kHz | 83.3 kHz | 30 kHz |  | 1.6 s |
| MAS: 13 kHz | sample 4 | <sup>13</sup>C: 55 kHz |  | 2500/15 ms | 200/6.6 ms |  | 42 h |
| H-H-C NOESY | 3D | HMQC type | <sup>1</sup>H-<sup>1</sup>H NOESY, 400 ms | <sup>1</sup>H | <sup>1</sup>H | <sup>13</sup>C | 12 |
| solution-state | 18.8 | delay 4 ms |  | 8 kHz | 1.5 kHz | 3.6 kHz | 1.6 s |
| NOESY-HMQC | sample 7 |  |  | 560/70 ms | 32/20 ms | 72/20 ms | 64 h |
| H-C-C NOESY | 3D | HMQC type | <sup>1</sup>H-<sup>1</sup>H NOESY, 350 ms | <sup>1</sup>H | <sup>13</sup>C | <sup>13</sup>C | 4 |
| solution-state | 22.3 | delay 4 ms |  | 11.4 kHz | 4.8 kHz | 4.8 kHz | 1.65 s |
| HMQC-NOESY-HMQC | samples 8-10, see Figure S5E |  |  | 571/50 ms | 95/20 ms | 70/14.7 ms | 60 h |

**Table S6.**
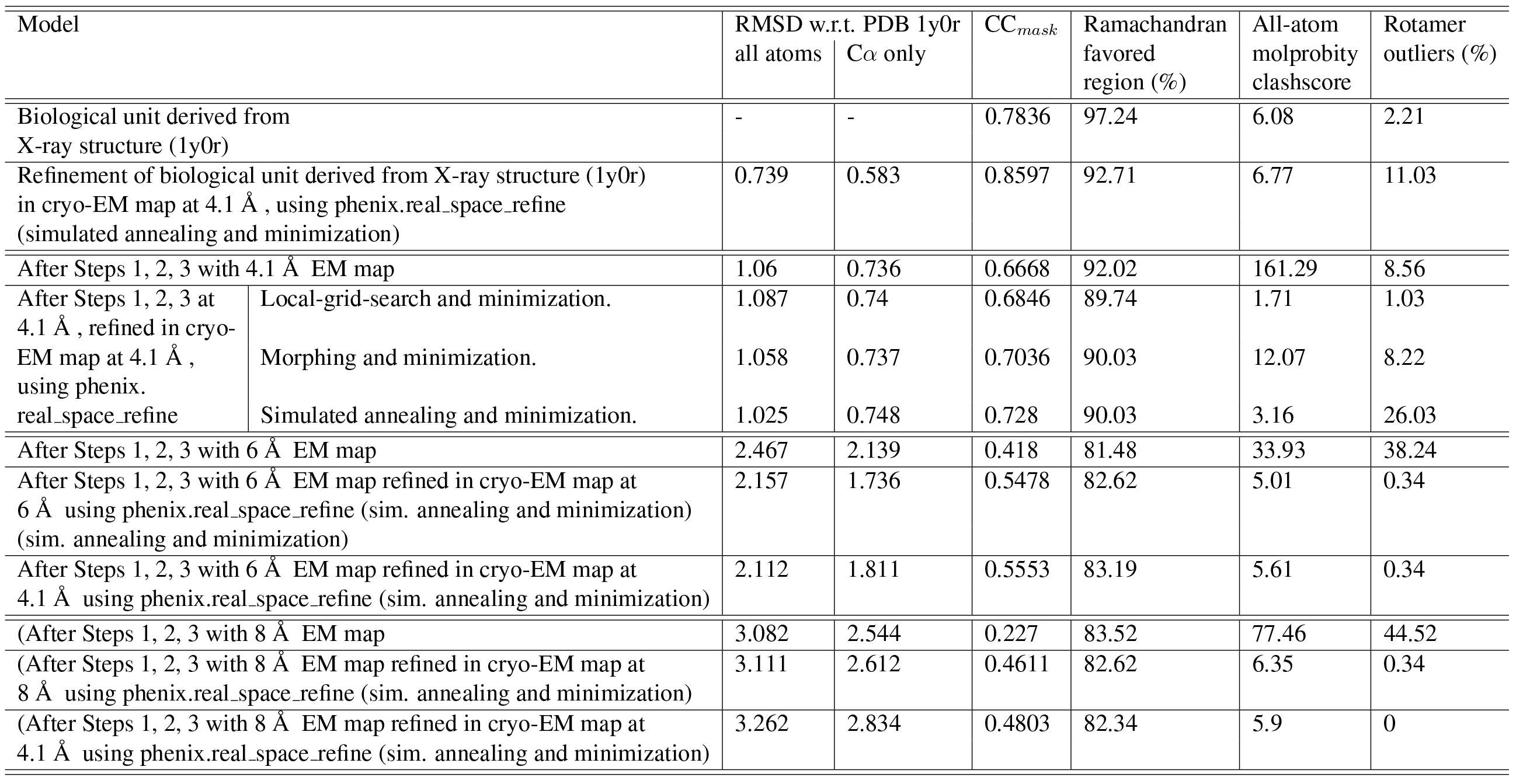
Refinement of TET2 structures with *phenix.real space-refine*. Shown are structure statistics obtained when using the three different EM maps (at 4.1, 6 or 8 Å resolution, respectively), and using as starting models different structures, either the crystal structure, or the structures obtained from the NMR and EM approach, that used 4.1, 6 or 8 Å resolution EM maps, respectively.

